# Pathogen protein modularity enables elaborate mimicry of a host phosphatase

**DOI:** 10.1101/2023.05.05.539533

**Authors:** Hui Li, Jinlong Wang, Tung Ariel Kuan, Bozeng Tang, Li Feng, Jiuyu Wang, Zhi Cheng, Jan Skłenar, Paul Derbyshire, Michelle Hulin, Yufei Li, Yi Zhai, Yingnan Hou, Frank L.H. Menke, Yanli Wang, Wenbo Ma

**Author notes:** Corresponding authors: Wenbo Ma or Yanli Wang. These authors contributed equally to this work (Hui Li, Jinlong Wang).

## Abstract

Pathogens produce diverse effector proteins to manipulate host cellular processes. However, how functional diversity is generated in an effector repertoire is poorly understood. Many effectors in the devastating plant pathogen *Phytophthora* contain tandem repeats of the “(L)WY” motif, which are structurally conserved but variable in sequences. Here, we discovered a functional module formed by a specific (L)WY-LWY combination in multiple *Phytophthora* effectors, which efficiently recruit the Serine/Threonine protein phosphatase 2A (PP2A) core enzyme in plant hosts. Crystal structure of an effector-PP2A complex shows that the (L)WY-LWY module enables hijacking of the host PP2A core enzyme to form functional holoenzymes. While sharing the PP2A-interacting module at the amino terminus, these effectors possess divergent C-terminal LWY units and regulate distinct sets of phosphoproteins in the host. Our results highlight the appropriation of an essential host phosphatase through molecular mimicry by pathogens and diversification promoted by protein modularity in an effector repertoire.

## INTRODUCTION

*Phytophthora* species, such as the Irish potato famine pathogen *Phytophthora infestans*, are major threats of global plant health and food security (Kamoun et al., 2015). Each *Phytophthora* genome encodes hundreds of effectors that are essential for disease development (Haas et al., 2009; Jiang et al., 2008; Wang and Jiao, 2019). The effector repertoires exhibit a high degree of diversity in different species, reflecting an accelerated evolution during host adaptation – a hallmark of co-evolutionary arms race (Dong and Ma, 2021; Wang and Jiao, 2019). It has been speculated that the (L)WY motif, named by conserved residues including multiple leucine (L), one tryptophan (W), and one tyrosine (Y) (He et al., 2019; Jiang et al., 2008), may contribute to effector evolution in *Phytophthora*. Prediction of five *Phytophthora* genomes revealed approximately three hundred effectors that consist of tandem repeats of the (L)WY motif. Structural analysis of one such effector, *<u>P</u>hytophthora* <u>s</u>uppressor of <u>R</u>NA silencing 2 (PSR2) of the soybean pathogen *P. sojae*, showed that each (L)WY unit forms a nearly identical **a**-helical bundle and adjacent units are connected through a conserved mechanism that results in joint-like linkages (He et al., 2019). Intriguingly, these effectors are often chimeras of (L)WY units with distinct surface residues (He et al., 2019), indicating variation in their capacity in interaction with host target(s). However, whether (L)WY motifs can serve as functional modules in these effectors by directly mediating specific interaction with host targets and how the modular architecture of the LWY effectors may promote the evolvability of effector repertoire in *Phytophthora* remains unknown.

Here, we use PSR2 as a model to investigate the role of (L)WY units in virulence functions and (L)WY-based modularity in effector evolution. We found that PSR2 associates with the Serine/Threonine protein phosphatase 2A (PP2A) core enzyme in plant hosts and forms a functional holoenzyme. By hijacking this host phosphatase, PSR2 alters the phosphorylation of specific host proteins to promote disease. Crystal structure and biochemical analysis of the PSR2-PP2A protein complex revealed (L)WY2-LWY3 of PSR2 to be responsible for competitive recruitment of the PP2A core enzyme. Structure-based search in two *Phytophthora* species identified 12 additional effectors that harbour this functional module and form effector-PP2A holoenzymes. Importantly, this PP2A-interacting module is always located on the amino terminus of the effectors, which possess divergent C-terminal LWY units. Using quantitative phosphoproteomics, we observed distinct impact of host proteins by two effectors, indicating that combining the PP2A-interacting module with different LWY units leads to functional diversification. This work highlights a major protein phosphatase as a key susceptibility target of *Phytophthora* pathogens and provides insight into how protein modularity may promote the diversity in an effector repertoire.

## RESULTS

### PSR2 hijacks PP2A core enzyme in plant hosts to promote disease

To investigate whether LWY units can mediate interactions with specific host molecules, we determined the interacting proteins of PSR2 in *Arabidopsis thaliana* using Immunoprecipitation followed by Mass Spectrometry (IP-MS). The results showed that PSR2 associated with the protein phosphatase 2A (PP2A) (**Figure 1A**). PP2A is responsible for the vast majority of Serine/Threonine phosphatase activity in eukaryotes and regulates diverse cellular functions (Brautigan, 2013; Fianu et al., 2021; Huang et al., 2020; Vervoort et al., 2021; Zheng et al., 2020). The highly conserved heterodimeric PP2A core enzyme, consisting of a scaffolding A subunit and a catalytic C subunit, is recruited by a regulatory B subunit to specific subcellular localization(s) and substrates (Brautigan, 2013; O’Connor et al., 2018). As such, the B subunit determines the function of the holoenzyme (O’Connor et al., 2018). Arabidopsis encodes three A subunits (RCN1, PDF1 and PDF2), five C subunits (PP2A-1 to PP2A-5) and 17 B subunits (Bian et al., 2020). PSR2 was co-precipitated with all the A and C subunits, but none of the B subunits (**Figures 1A and 1B**). Importantly, PSR2 protein complex enriched from the transgenic plant tissue possessed a phosphatase activity that could be completely inhibited by the PP2A-specific inhibitor okadaic acid (**Figure 1C**), suggesting that PSR2 forms a functional holoenzyme with the Arabidopsis PP2A core enzyme in planta. The interaction of PSR2 with PP2A core enzyme and the formation of a functional holoenzyme was also confirmed in *Nicotiana benthamiana* expressing PSR2 (**Figure S1A and S1B**).

**Figure 1.**
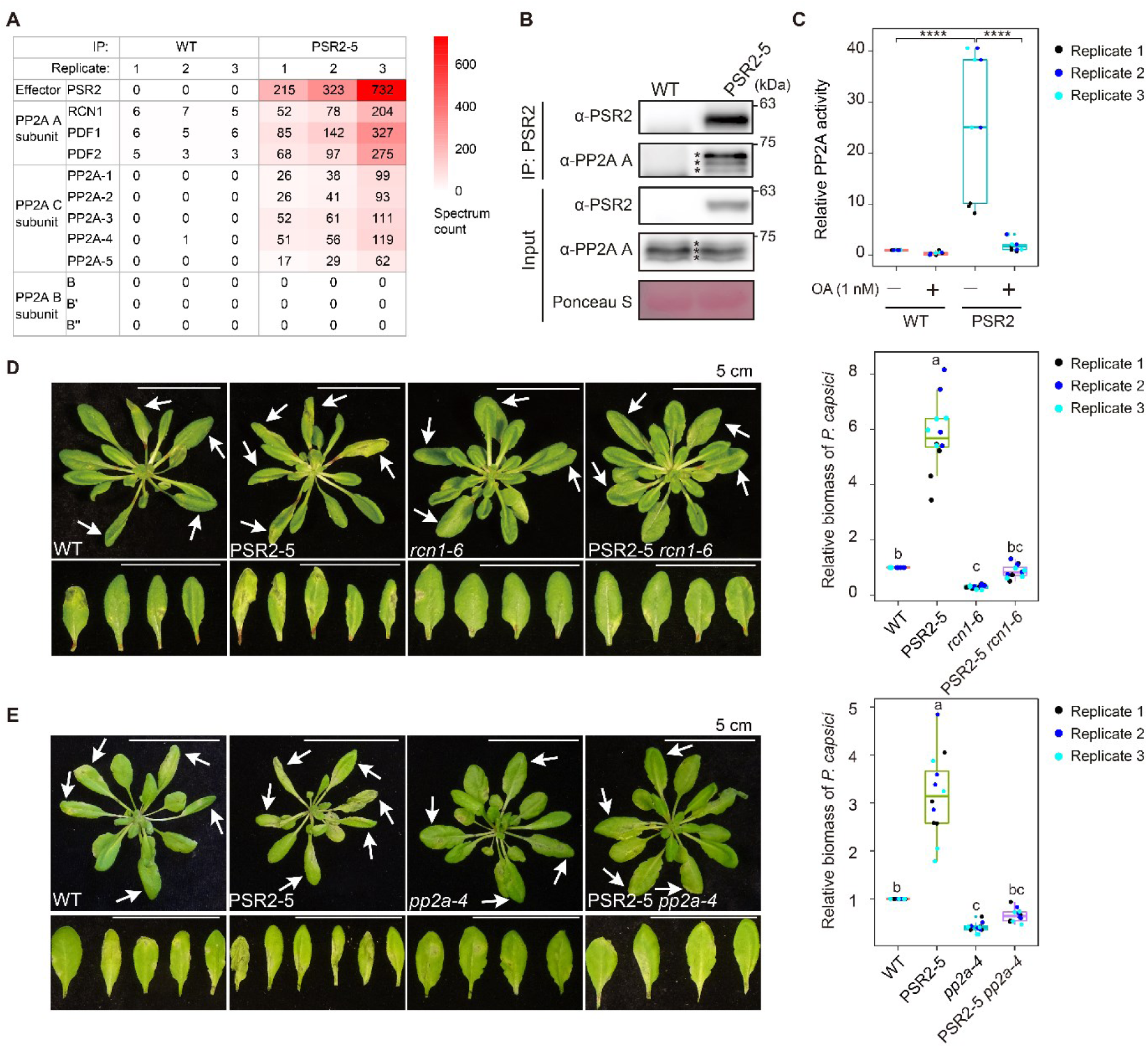
The Ser/Thr protein phosphatase PP2A core enzyme is a susceptibility factor targeted by the *Phytophthora* effector PSR2. (A) PSR2 associates with PP2A A and C subunits, but not B subunits in Arabidopsis. Numbers of spectrum counts detected by IP-MS are presented. WT, *Arabidopsis thaliana* ecotype Col-0; PSR2-5, transgenic Arabidopsis expressing PSR2 in Col-0 background. (B) PSR2 co-immunoprecipitated with PP2A A subunits. PSR2 was enriched from protein extract of 2-week-old Arabidopsis seedlings using an anti-PSR2 antibody. PDF1, RCN1 and PDF2 (labelled by asterisks from top to bottom) were detected using an anti-PP2A A antibody by western blotting. Ponceau S staining was used to assess equal loading. The interaction between PSR2 and PP2A A subunits in *N. benthamiana* is presented in Figure S1A. (C) PSR2 complex possessed PP2A phosphatase activity. Protein complexes enriched by an anti-PSR2 antibody as described in (B) were examined for phosphatase activity using a phosphopeptide as the substrate. Okadaic acid (OA) is a PP2A specific inhibitor. Values from three independent experiments were analyzed by two-tailed Student t-test (\*\*\*\**p* < 0.0001). The PP2A phosphatase activity of PSR2 complex in *N. benthamiana* is shown in Figure S1B. (D and E) Virulence activity of PSR2 requires the PP2A A subunit RCN1 (D) and PP2A C subunit PP2A-4 (E). Four-week-old Arabidopsis plants were inoculated with *Phytophthora capsici* isolate LT263. Photos were taken at 3 days post inoculation (dpi) with arrows indicating inoculated leaves. Relative biomass of *P. capsici* in inoculated plants at 3 dpi was determined (n ≥ 20 in each sample per experiment) and data from three independent replicates are presented. One-way ANOVA and post hoc Tukey were used for statistical analysis. Different letters label significant differences (*p* < 0.05). *P* values for all the experiments are provided in Table S6.

PSR2 was reported to increase Arabidopsis susceptibility to *Phytophthora capsici* (Hou et al., 2019; Qiao et al., 2013). We expressed PSR2 in the PP2A A subunit mutant *rcn1* and observed that this virulence activity was abolished (**Figure 1D; Figure S1C**). The virulence activity of PSR2 was also compromised in another PP2A A subunit mutant *pdf1* (**Figure S1C-S1F**), indicating that the PP2A A subunits are required for PSR2 to promote disease. Since the PP2A enzymatic activity is determined by the catalytic C subunit, we generated transgenic Arabidopsis expressing PSR2 in the C subunit mutant *pp2a-4*. Similar to *rcn1*, the virulence activity of PSR2 was also abolished in the *pp2a-4* background (**Figure 1E; Figure S1C**), suggesting that the cellular function of PSR2 in plant hosts depends on the PP2A phosphatase activity. It is important to note that none of the PP2A A or C subunit mutants, including higher order mutants, was hypersusceptible to *P. capsici* (**Figure S1G and S1H**). Taken together, these results suggest that the PSR2 virulence function could not be attributed to a reduced PP2A activity in Arabidopsis. Indeed, the *rcn1* and *pp2a-4* mutants showed enhanced resistance to *P. capsici* (**Figure 1D and 1E; Figure S1G and S1H**), consistent with their role as a susceptibility target of PSR2.

### PSR2 forms a complex with PP2A A subunit through LWY2-LWY3

To determine whether PSR2 can directly interact with Arabidopsis PP2A A and/or C subunits in the core enzyme, we carried out *in vitro* pull-down and gel filtration assays. Despite numerous trials using different expression systems, we were unable to obtain soluble proteins of the C subunits, which were in inclusion bodies. Therefore, a highly conserved human C subunit (C**a**) (**Figure S2A**) was used as an alternative. Our results show that PSR2 forms stable binary complexes with both RCN1 and PDF1 (**Figure 2A; Figure S2B-S2D**), but not with C**a** (**Figure S2E**).

**Figure 2.**
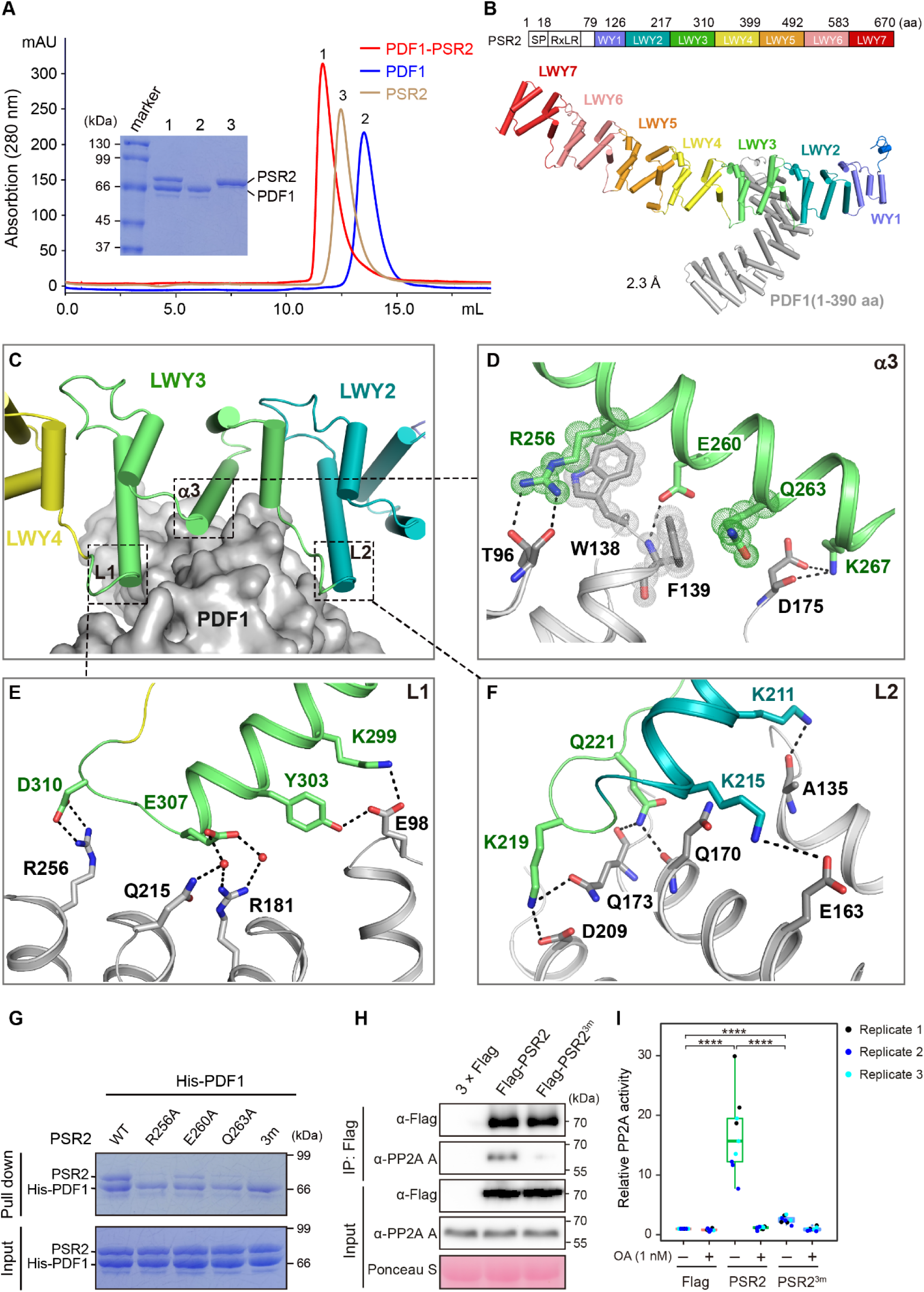
PSR2 interacts with a PP2A A subunit through an interface formed by (L)WY2-LWY3. (A) Gel filtration chromatography shows the formation of a PDF1-PSR2 binary complex. The three peaks represent PDF1-PSR2 complex, apo state PDF1, and apo state PSR2, respectively. The formation of PSR2 binary complex with a truncated PDF1(1-390 aa) is shown in Figure S2D. (B) Crystal structure of the binary PSR2(59-670aa)-PDF1(1-390 aa) complex with a resolution of 2.3 Å. The WY1-(LWY)_6_ arrangement of PSR2 is also presented. SP, secretion signal peptide; RXLR, a translocation motif of *Phytophthora* effectors. (C) The PSR2-PDF1 interaction interface in centered on the third α-helix (α3) of LWY3 (lime) and strengthened by L1 and L2 on both sides. (D-F) Twelve residues in α3 (D), L1 (E) and L2 (F) directly interact with PDF1 (grey). (G) *In vitro* pull-down assay confirms the REQ triad (R256, E260, Q263) in the α3 of LWY3 is essential for PDF1 interaction. 3m: PSR2^R256A/E260A/Q263A^. Pull down results for each individual PSR2 mutants with PDF1 are shown in Figure S2I. (H and I) The REQ triad is required for forming a functional PSR2-PP2A holoenzyme *in planta*. Flag-tagged PSR2 or PSR2^3m^ were expressed in *Nicotiana benthamiana* and immunoprecipitated using anti-Flag magnetic beads. Co-immunoprecipitation of PP2A A subunit(s) was detected using an anti-PP2A A antibody (H) and phosphatase activity was measured with or without the PP2A inhibitor Okadaic acid (OA) (I). Values from three independent repeats were analyzed by two-tailed Student t-test (\*\*\*\**p* < 0.0001). *P* values for all the experiments are provided in Table S6.

To further understand the molecular details of the direct interaction of PSR2 with PP2A A subunits, PDF1-PSR2 and RCN1-PSR2 complexes were subjected to extensive screening of crystallization conditions. We eventually solved the crystal structure of the PSR2(59-670 aa)-PDF1(1-390 aa) complex at 2.3 Å (**Figure 2B; Table S1**). In this complex, PDF1 exhibits an arc-like structure with its N-terminal portion interacting with the LWY2-LWY3 region of PSR2, which has a stick-like WY1-(LWY)_6_ overall architecture as described previously (He et al., 2019). The WY1 and LWY4-LWY7 units of PSR2 are located on the outer and inner sides of the PDF1 arc, respectively. PSR2 forms an extensive interaction interface with the third α-helix of LWY3 (thereafter α3) serving as a core. α3 is embedded in a small groove on PDF1 surface, forming multiple hydrogen bonds and **p**-**p** interactions with PDF1 (**Figure 2C and 2D**). In particular, the residues R256 and Q263 within α3 stack on the side-chains of PDF1^W138^ and PDF1^F139^, respectively, strengthening the complex by **p**-**p** interactions (**Figure 2D**). Furthermore, R256 and E260 form hydrogen bonds with PDF1^T96^ and PDF1^F139^ respectively. Thus, this “REQ” triad of α3 plays a central role in the PSR2-PDF1 complex formation. Linkages on both sides of LWY3 (hereafter L1 and L2, **Figure 2C**) are rich of charged residues, including K299, Y303, E307 and D310 in L1 (**Figure 2E**), and K211, K215, K219 and Q221 in L2 (**Figure 2F**). These residues further stabilize the complex by forming salt bridges or hydrogen bonds with PDF1. Upon binding to PDF1, PSR2 undergoes a significant conformational change in which L1 and L2 move towards each other to capture PDF1 (**Figure S2F-S2H**). Together, the PSR2-PDF1 complex structure revealed that the α3 of LWY3 serves as an interaction center to hold PDF1 in place, and L1 and L2 act like tweezers to stabilize the complex. Considering that PDF1 and RCN1 share 96% sequence similarity, the same mechanism is likely involved in PSR2 interaction with RCN1. Indeed, 11 of the 12 PSR2-interacting residues are conserved in RCN1 or PDF2 (data not shown).

To confirm the interactions identified in the PSR2-PDF1 complex, we mutated 11 residues within the L1-α3-L2 region of PSR2 and examined their impact on interaction with PDF1 *in vitro*. The results show that the interaction was decreased in many mutants including PSR2^R256A^ and PSR2^Q263A^ in the REQ triad (**Figure S2I**). Consistent with the role of this triad as the center of the interaction interface, the mutant PSR2^R256A/E260A/Q263A^ was nearly abolished for PDF1 interaction *in vitro* (**Figure 2G**) and *in planta* (**Figure 2H**). In addition, phosphatase activity was no longer detectable in the PSR2^R256A/E260A/Q263A^ protein complex when the mutant was expressed in *N. benthamiana* (**Figure 2I**). These results confirmed the interaction interface defined by the PSR2-PDF1 complex structure, and more importantly, allowed us to assign a specific function, i.e. mediating direct interactions with PDF1 or likely other PP2A A subunits in the hosts, to the LWY2-LWY3 units of PSR2 (**Figure S2J**), demonstrating that (L)WY units can serve as functional modules.

### PSR2 binds tighter to the PP2A core enzyme than an Arabidopsis B subunit

The structure of PDF1 in the PSR2-PDF1 complex shows a high level of conservation with the human PP2A A subunit PPP2R1A (Groves et al., 1999) (**Figure S3A**). Interestingly, although PSR2 is structurally distinct from all the known human PP2A B subunits, they bind to a similar region on the N-terminal portion of the A subunit (Cho and Xu, 2007; Wlodarchak et al., 2013; Xu et al., 2008; Xu et al., 2006) (**Figure 3A**). Indeed, several residues conserved in PDF1 and PPP2R1A are involved in the interactions with PSR2 and human B subunits respectively (**Figure 3B; Figure S3B**). For example, PDF1^W138^ forms a critical **p**-**p** interaction with PSR2^R256^; its counterpart PPP2R1A^W140^ is also involved in the interactions with human B subunits that belong to three different subfamilies (**Figure 3B; Figure S3B**). These results suggest that the binding site in PDF1 for PSR2 likely overlaps with the site that binds the endogenous plant B subunits, leading to their mutual exclusion in PP2A holoenzymes. This is consistent with our IP-MS results, which showed that none of the Arabidopsis PP2A B subunits was associated with PSR2 (**Figure 1A**). Together with the observation that PSR2 protein complex possesses PP2A phosphatase activity, these findings support that PSR2 functions as a pathogen-derived PP2A B subunit in plant cells.

**Figure 3.**
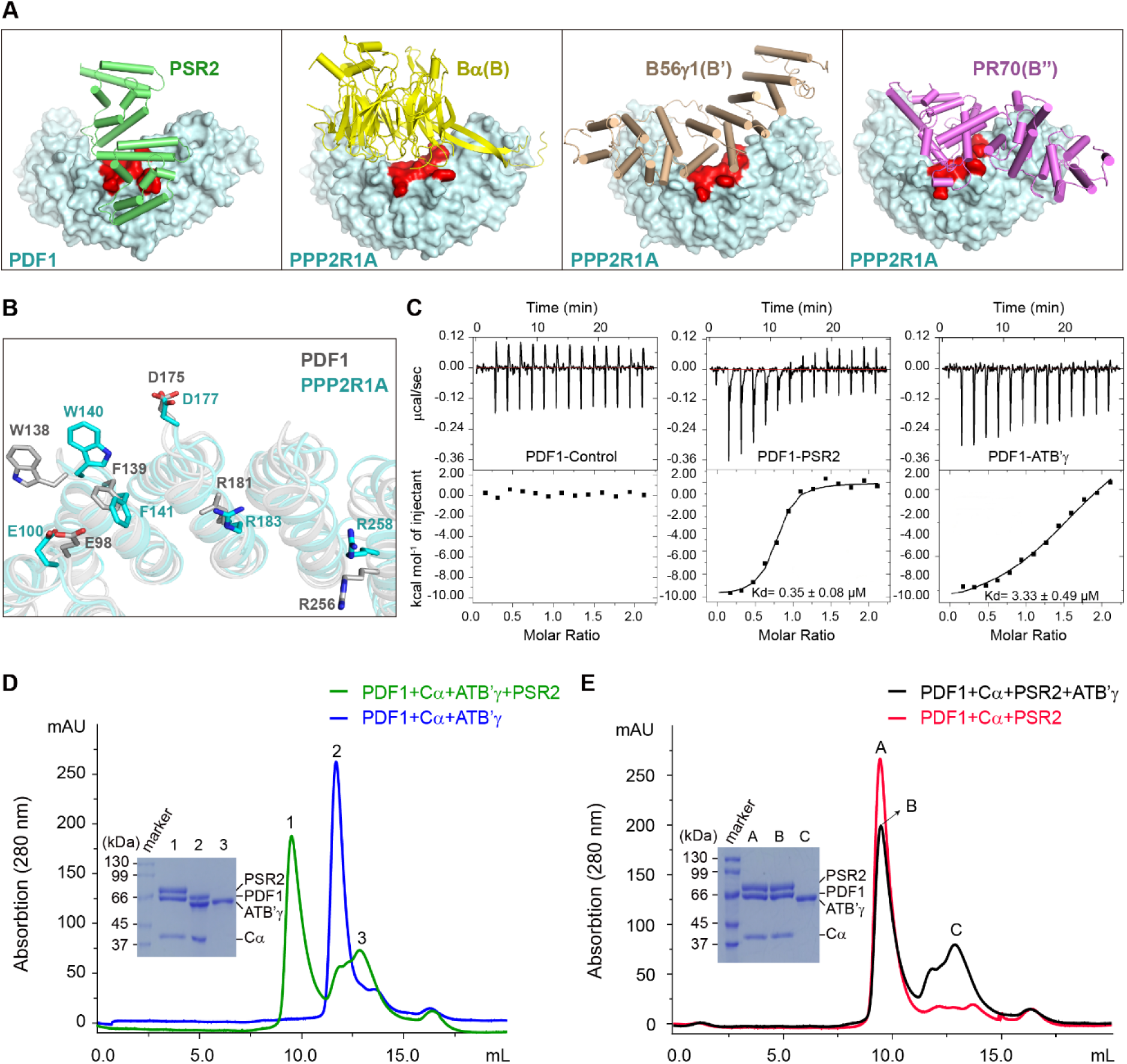
PSR2 is a molecular mimic of PP2A B subunit that can efficiently recruit host core enzyme to form a functional holoenzyme. (A) The PSR2-binding site in PDF1 (in red) is conserved in human PP2A A subunit PPP2R1A to bind endogenous B subunits. B**a**, B56**g** and PR70 belong to the B, B’, and B’’ subfamilies, respectively. Structural alignment of PDF1 and PPP2R1A is shown in Figure S3A. (B) Conserved residues in PDF1 (grey) and PPP2R1A (cyan) interact with PSR2 and the human B subunits respectively. Sequence alignment of PDF1 (1-390 aa) and PPP2R1A (1-396 aa) with residues involved in the interaction with PSR2 and endogenous human B subunits are shown in Figure S3B. (C) Isothermal Titration Calorimetry (ITC) measurement of binding affinities of PDF1 with PSR2 vs an Arabidopsis PP2A B subunit ATB’**g**. (D) PSR2 efficiently displaced ATB’**g** from a preformed PDF1-ATB’**g**-C**a** holoenzyme *in vitro*. Gel filtration chromatography (in green) shows two peaks representing the PDF1-PSR2-C**a** holoenzyme and excess ATB’**g** dropped from the preformed PDF1-ATB’**g**-C**a** holoenzyme. Chromatography of the PDF1-ATB’**g**-C**a** holoenzyme (in blue) was used as a control. (E) ATB’**g** could not efficiently displace PSR2 in a preformed PDF1-PSR2-C**a** holoenzyme. Gel filtration chromatography (in black) shows the PDF1-PSR2-C**a** holoenzyme and ATB’**g** added to the mixture. Chromatography of the PDF1-PSR2-C**a** holoenzyme (in red) was used as a control.

To examine whether PSR2 could efficiently compete with endogenous B subunits when recruiting the PP2A core enzyme, we compared the binding affinity of PDF1 with PSR2 or an Arabidopsis PP2A B subunit ATB’**g** (Trotta et al., 2011). Using isothermal titration calorimetry (ITC), the K_d_ for PSR2-PDF1 binding was estimated to be 0.35 ± 0.08 µM. In comparison, the K_d_ for ATB’**g**-PDF1 binding was approximately 3.33 ± 0.49 µM, ∼10-fold higher than PSR2-PDF1 (**Figure 3C**). To directly examine whether PSR2 binds tighter to the PP2A core enzyme than ATB’**g**, we conducted an *in vitro* competition assay using a heterodimer consisting of PDF1 and the human PP2A C subunit C**a**. When adding to a preformed PDF1-C**a**-ATB’**g** complex, PSR2 facilitated the displacement of ATB’**g** and formed its own holoenzyme with the PDF1-C**a** core enzyme (**Figure 3D**). In contrast, ATB’**g** couldn’t efficiently dissociate PSR2 from a preformed PDF1-C**a**-PSR2 holoenzyme (**Figure 3E**). These results demonstrate that PSR2 can competitively recruit the PP2A core enzyme as a molecular mimic of B subunits.

### The PP2A-interacting module is adopted in multiple LWY effectors

Structural characterization of the PSR2-PDF1 complex defined a PP2A-interacting module that involves ten residues in LWY3 and two residues in the “WY” portion of LWY2 (**Figure S2J**). Considering the modular architecture of the LWY effectors, we reasoned that any effectors harbouring the PSR2 (L)WY2-LWY3 module would also be able to recruit the PP2A core enzyme. To test this, 134 WY1-(LWY)n effectors predicted from *P. sojae* and *P. infestans* (He et al., 2019) were subjected to a similarity search based on both structure and sequence. These effectors were first analyzed by Alphafold2 (Jumper et al., 2021) (**Document S1**); then all the (L)WY-LWY pairs defined in the structural models were screened for structural and sequence similarity to the PP2A-interacting module in PSR2 (**Table S2**). Candidates were further examined for conservation at the positions corresponding to the 12 PDF1-interacting residues (**Table S2**). Based on the results, 15 effectors were further tested by experimental confirmation. These effectors were individually expressed in *N. benthamiana* to enrich the effector protein complexes, from which their interaction with the PP2A A subunits and possession of phosphatase activity were examined (**Figure 4A; Figure S4A-S4C**). An additional four effectors were included in the experimentation as negative controls. The results show that 12 of the 15 candidate effectors, including a PSR2 ortholog in *P. infestans* (named PiPSR2), formed functional PP2A holoenzymes *in planta* (**Figure 4A; Figure S4A-S4C**). Thus, there is a strong corelation between the possession of a PSR2-like PP2A-interacting module and the ability to hijack host PP2A core enzyme by *Phytophthora* effectors. From these 13 LWY effectors that can associate with the plant PP2A core enzyme, we made the observation that they all contain a minimum of five WY/LWY units and the predicted PP2A-interacting module is always located at the amino terminus (**Figure 4A**). These findings indicate that a similar mechanism is employed by these effectors for their association with PP2A core enzyme.

**Figure 4.**
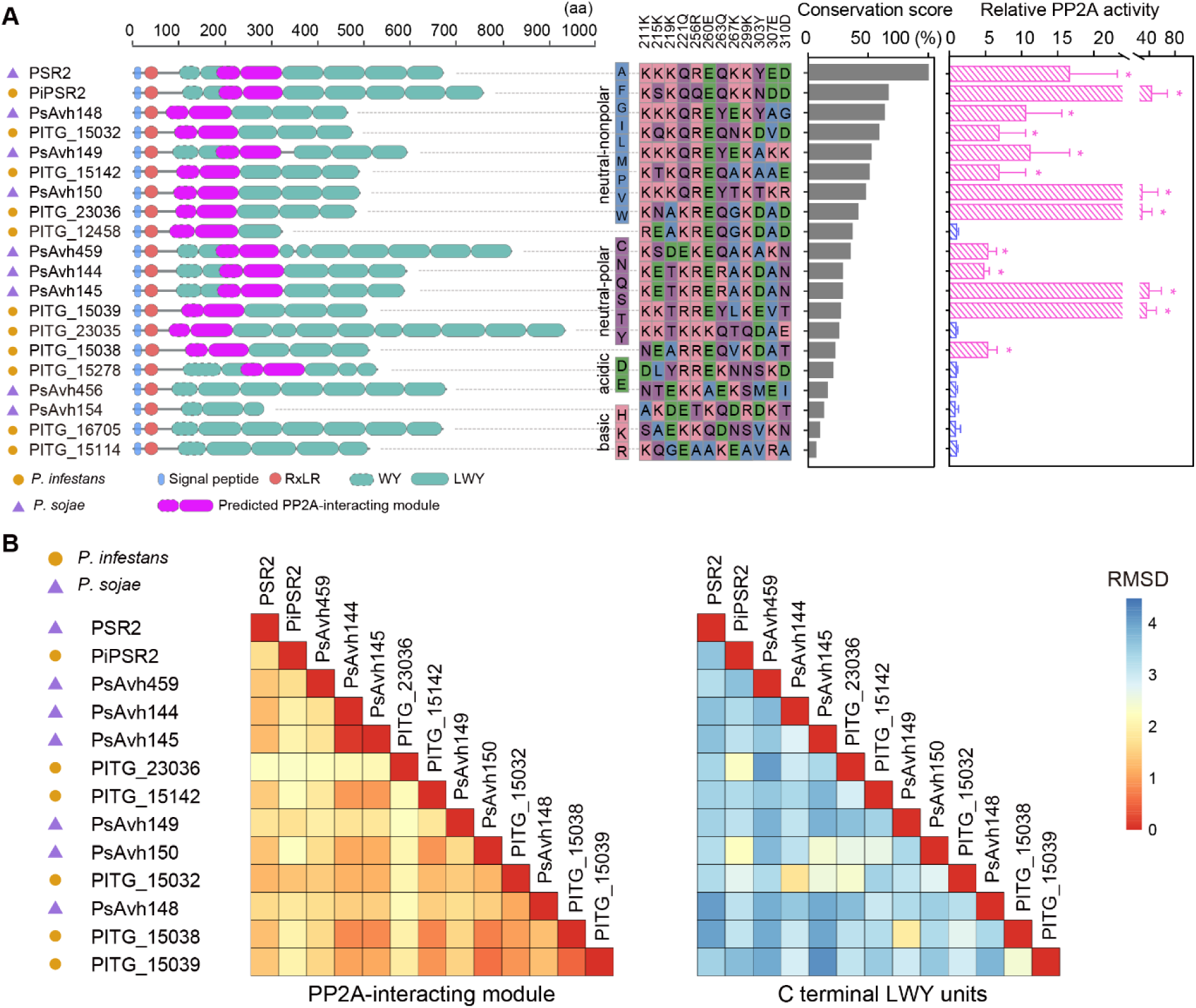
Twelve effectors adopted the PSR2-like, PP2A-interacting module at the N-terminus but have diverse C-terminal LWY units. (A) Twelve LWY effectors predicted to possess the PP2A-interacting module can form functional PP2A holoenzymes. Effector architecture showing the LWY unit arrangement and the conservation of the 12 PDF1-interacting residues is presented for 15 candidates and four negative controls. The predicted PP2A-interacting modules are highlighted in pink. The conservation score is a sum value predicted for each residue based on structural superimposition and BLOSUM62 matrix. Predicted structure models of the effectors are shown in Supplementary Data 1 and detailed information on conservation scores is shown in Table S2. These effectors were individually expressed in *N. benthamiana* and examined for PP2A phosphatase activity. Values from three independent experiments were analyzed by two-tailed Student t-test (*p* < 0.01). Experimental results of effector interaction with PP2A A subunits and the phosphatase activity of each effector complex are shown in Figure S4A-S4C. *P* values for all the experiments are provided in Table S6. (B) C-terminal LWY units exhibit a higher level of diversity in the PP2A-associating effectors. RMSD values from pairwise analysis are used to reflect structural similarity.

### Effector-PP2A holoenzymes are diverse

Substrate specificity of a PP2A holoenzyme is determined by the specific B subunit (Janssens and Goris, 2001; Shi, 2009). Previous research of human PP2A holoenzymes suggests that regions on the regulatory B subunits located close to the PP2A C subunit in the complex are important for substrate recruitment. It is intriguing that *Phytophthora* produces a suite of effectors that all function as molecular mimics of B subunits by adopting the same PP2A A-interacting module, which is always located on the N terminus. Similarity in the structural organization between the PSR2-PDF1 complex and human PP2A holoenzymes indicates that the PP2A C subunit would be located close to the C-terminal LWY units of PSR2. Therefore, the C-terminal LWY units might be responsible for substrate binding. As such, divergence in the C-terminal LWY units of the PP2A-interacting effectors may lead to functional diversification. To investigate the potential diversity in the function of PP2A-interacting effectors, we analyzed the evolutionary trajectories of C- vs N-terminal LWY units in the PP2A-recruiting effectors. Intriguingly, pairwise analysis of structural and sequence similarity revealed an overall higher diversity at the C-terminal LWY units compared to the PP2A-interacting units on the N terminus (**Figure 4B; Table S3; Figure S4D**). We also determined the subcellular localization of the PP2A-interacting effectors in plant cell by transient expression in *N. benthamiana*. Confocal microscopy showed diverse localization patterns of the effectors (**Figure S4E**), which would also contribute to a potential divergence in substrate-binding capacity of the effector-PP2A holoenzymes.

### PSR2 and PITG_15142 regulate the phosphorylation of different host proteins

To further investigate the functional conservation and diversity of the PP2A-interacting effectors, we determined the crystal structure of the *P. infestans* effector PITG_15142 at 3.1 Å (**Figure 5A; Table S1**). PITG_15142 has a WY1-(LWY)_4_ architecture with the predicted PP2A-interacting module residing in the very N-terminal (L)WY units. Indeed, WY1-LWY2 of PITG_15142 has a significant structural similarity with (L)WY2-LWY3 of PSR2 with the root-mean-square deviation (RMSD) score of 1.38 Å (**Figure 5B**). Seven residues in PSR2 that mediate direct interactions with PDF1 are conserved in PITG_15142. These conserved residues include R167, E171, and Q174, which correspond to the REQ triad (**Figure 5B, Figure S5A**). Similar to PSR2, mutation of the REQ triad abolished the interaction of PITG_15142 with PDF1 *in vitro* (**Figure 5C**) and *in planta* (**Figure S5B**). In addition, phosphatase activity was no longer detectable in the PITG_15142^R167A/E171A/Q174A^ protein complex when the mutant was expressed in *N. benthamiana* (**Figure S5C**). The PITG_15142^R167A/E171A/Q174A^ mutant was also unable to promote *P. capsici* infection in *N. benthamiana* (**Figure 5D; Figure S5D**), suggesting that the REQ triad of PITG_15142 is essential for its functions in plant cells. Furthermore, PITG_15142 formed a stable ternary complex with the core PP2A enzyme PDF1-C**a** *in vitro* (**Figure S5E**). Together, these results validated that PITG_15142, and possibly the other LWY effectors possessing the PP2A-interacting module, interacts with PDF1 and forms an effector-PP2A holoenzyme through the same mechanism employed by PSR2.

**Figure 5.**
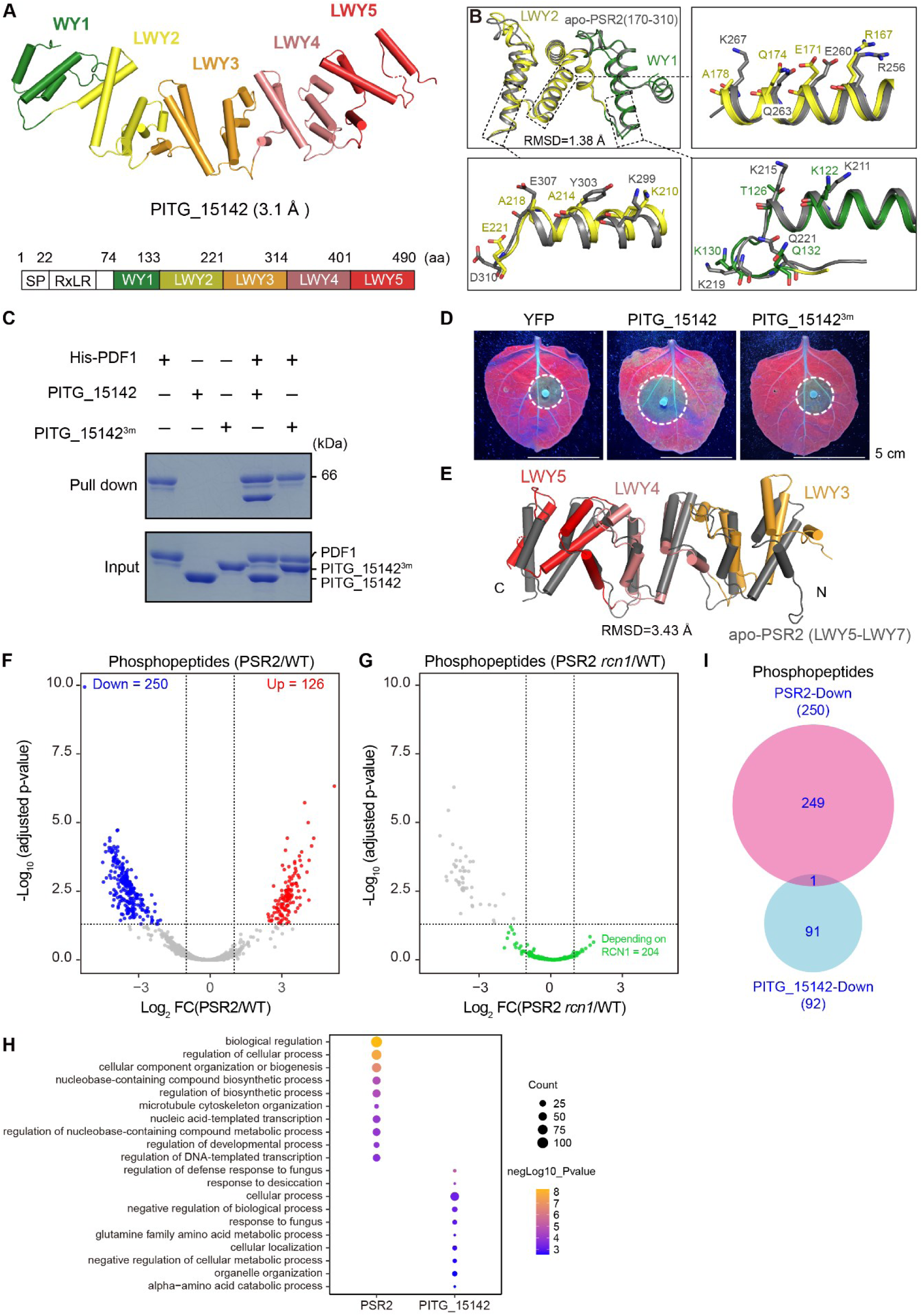
PSR2 and PITG_15142 affect the phosphorylation of distinct sets of host proteins. (A) Crystal structure of the *P. infestans* effector PITG_15142 with a resolution of 3.1 Å. PITG_15142 has a WY1-(LWY)_4_ architecture with the predicted PP2A-interacting module located at WY1-LWY2. (B) Structural superimposition of the PP2A-interacting module in PITG_15142 (81-221 aa, in forest for WY1 and tv_yellow for LWY2) and the apo state PSR2 (170-310 aa, in grey) shows a high similarity (RMSD=1.38 Å) and seven of the 12 residues that mediate PSR2 interaction with PDF1 are conserved in PITG_15142. In particular, the α3 of PSR2, which serves as the interaction core with PDF1, is highly conserved in LWY2 of PITG_15142 Sequence alignment of PITG_15142 and PSR2 with key residues involved in the interaction with PDF1 is shown in Figure S5A. (C) *In vitro* pull-down showing direct interaction of PITG_15142 with PDF1, which requires the REQ triad. PITG_15142^3m^ contains the R167A/E171A/Q174A mutations. Gel filtration chromatography showing one single peak representing the PDF1-PITG_15142-C**a** holoenzyme is presented in Figure S5B. (D) The REQ triad is required for the virulence activity of PITG_15142 in *N. benthamiana*. PITG_15142 or PITG_15142^3m^ were expressed in *N. benthamiana* using *Agrobacterium*-mediated transient expression. 48 hours after Agro-infiltration, the leaves were inoculated with zoospore suspensions of *P. capsici*. Photos were taken at 3 days post inoculation. Quantitative analysis of the lesion areas is presented in Figure S5D. (E) PSR2 and PITG_15142 have divergent C-terminal units. Structural superimposition of the C-terminal units in PSR2 LWY5-LWY7 (400-670 aa) and PITG_15142 LWY3-LWY5 (222-490 aa) shows a RMSD value of 3.43 Å. (F) PSR2 affects phosphoproteome of Arabidopsis. Volcano plot shows changes of phosphopeptide abundance according to the average ratio (log_2_) and *p* value (-log_10_ adjusted *p*-value) in Arabidopsis plants expressing PSR2. Gray dots represent phosphopeptides with non-significant change in abundance. Red and blue dots represent phosphopeptides with significantly increased or decreased abundance, respectively. (G) Volcano plot shows changes of the PSR2-reduced phosphopeptides in plants expressing PSR2 in the *rcn1-6* background. 204 of the 250 phosphopeptides (shown as green dots) were no longer reduced in their relative abundance levels in the *rcn1-6* background. The vertical dashed lines indicate *p*-value = 0.05 and the horizontal dashed line indicates a fold change of 2. (H) Gene ontology (GO) analysis of phosphoproteins with reduced phosphorylation levels in Arabidopsis expressing PSR2 or PITG_15142. The top 10 (*P* value <0.01, ranked by OddsRatio) significantly enriched GO terms in biological process group are displayed. (I) Venn diagram of significantly reduced phosphopeptides in Arabidopsis expressing PSR2 or PITG_15142.

Despite the similarity in PP2A interaction, a comparison of the PSR2 and PITG_15142 structures revealed divergence in the C-terminal units. A comparison of PSR2 (LWY5-LWY7) and PITG_15142 (LWY3-LWY5) revealed an RMSD value of 3.43 Å (**Figure 5E**), indicating a potential diversification in substrate binding. To determine whether PSR2 and PITG_15142 recruit different host proteins to the effector-PP2A holoenzymes, we generated transgenic Arabidopsis expressing PITG_15142 tagged with TurboID. The transgenic plants were hypersusceptible to *P. capsici* (**Figure S5F and S5G**), confirming that PITG_15142 has a virulence function in Arabidopsis. We then performed a label free quantitative phosphoproteomic analysis using liquid chromatography-mass spectrometry (LC-MS). In total, 10696 unique phosphopeptides from 4148 proteins were detected and quantified (**Table S4**). A comparison of the phosphoproteome profiles between PSR2-5 vs wild-type and PITG_15142-YFP-TurboID vs YFP-TurboID plants did not reveal a substantial overall change (**Table S4**), indicating that these effectors do not have a global impact on protein phosphorylation. However, they influenced the phosphorylation status of specific peptides. In PSR2-5, 250 peptides showed reduced phosphorylation and 126 showed increased phosphorylation (Fold change >2, adjusted *P* value <0.05, in four biological replicates) (**Figure 5F**). The phosphopeptides with reduced phosphorylation in PSR2-expressing plants are of particular interest as it may represent the phosphatase activity from the specific effector-PP2A holoenzyme. Importantly, 204 out of the 250 phosphopeptides that showed reduced phosphorylation in PSR2-5 were no longer altered when PSR2 was expressed in the *rcn1-6* mutant background **(Figure 5G)**. These results demonstrate that PSR2-mediated host protein dephosphorylation depends on Arabidopsis PP2A core enzyme. In PITG_15142-expressing plants, 92 and 159 peptides showed reduced or increased phosphorylation, respectively (**Figure S5H**). Gene Ontology (GO) analysis of proteins with reduced phosphorylation by PSR2 suggests enrichment in nucleobase-containing compound biosynthetic process (**Figure 5H**). However, peptides with reduced phosphorylation by PITG_15142 are mainly involved response to fungus and desiccation (**Figure 5H**). Importantly, only one peptide showed decreased phosphorylation in both PSR2- and PITG_15142-expressing plants, consistent with the notion that these two effectors regulate the phosphorylation of distinct sets of host proteins (**Figure 5I**).

To further confirm PSR2 and PITG_15142 have different binding capacities with host proteins, we determined the interacting proteins of PITG_15142 in Arabidopsis using IP-MS and then compared the interactome with that of PSR2. The only common interacting proteins between these two effectors are the PP2A core enzyme subunits (**Figure S5I; Table S5**). These results confirmed that effectors harbouring the PP2A-interacting module form holoenzymes with the host core enzyme but regulate different phosphoproteins, presumably based on their C-terminal LWY units. As such, these effectors exhibit functional diversity, which is enabled by the modular architecture based on LWY tandem repeats.

## DISCUSSION

Being the major protein phosphatase, PP2A regulates a large variety of cellular processes and its dysfunction is associated with many diseases (Brautigan, 2013; Fianu et al., 2021; Huang et al., 2020; Vervoort et al., 2021; Zheng et al., 2020). Human PP2A is a ubiquitous virulence target by oncogenic viruses, some of which can also bind to the core enzyme (Barski et al., 2021). However, these viral proteins mostly inhibit the PP2A enzymatic activity rather than mimicking the function of B subunits. In plants, PP2A has been reported to regulate immunity (Segonzac et al., 2014) and associate with the virulence function of the type III-secreted effectors AvrE of the bacterial pathogen *Pseudomonas syringae* and its homolog DspA/E in *Erwinia amylovora* (Jin et al., 2016; Siamer et al., 2014). In addition, the effector HaRxL23 produced by an oomycete pathogen *Hyaloperonospora arabidopsidis* can partially complement the virulence activity of AvrE, indicating that HaRxL23 may also manipulate host PP2A (Deb et al., 2018). Our identification of multiple PP2A B-mimicking effectors in *Phytophthora* established that PP2A is a key susceptibility factor widely targeted by pathogens across the kingdom through independent mechanisms. This is consistent with the resistance phenotype of PP2A core enzyme mutants to both *Pseudomonas syringae* (Segonzac et al., 2014) and *Phytophthora capsici*. Our detailed structural characterization of the effector-PP2A complex offers new opportunities to enhance disease resistance through precise engineering.

Accelerated effector evolution is essential for host adaptation of pathogens (Dong and Ma, 2021; Ma and Guttman, 2008; Outram et al., 2022). The elaborate host mimicry of a key enzyme complex by multiple *Phytophthora* effectors through the adoption of the same functional module demonstrate how protein modularity can facilitate the expansion of virulence targets in a predictable manner. The structurally conserved but functionally variable (L)WY units may serve as a reservoir of functional modules. Shuffling of these modules as tandem repeats can then enhance the evolvability of an effector repertoire through which new activities of host manipulation could arise (**Figure 6**). In bacterial pathogens, “reassortment” of DNA sequences spanning the promoter and N-terminal translocation signal was also proposed as a mechanism through which new Type III secreted effectors could be created (Stavrinides et al., 2008). Therefore, shuffling of functional units based on modular protein/gene architecture is a common theme in the evolution of pathogenicity.

**Figure 6.**
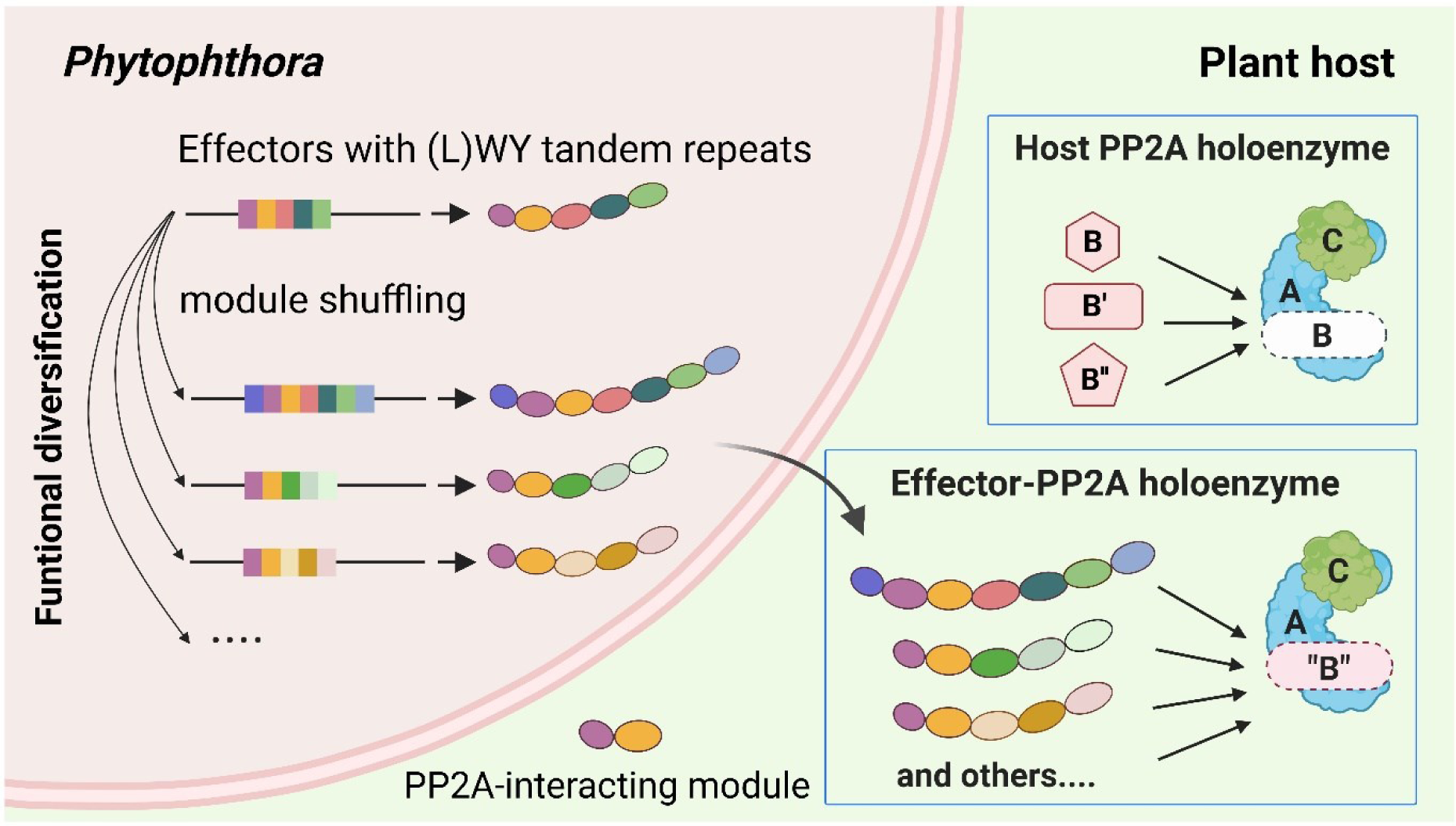
A model illustrating how effector evolution might be promoted by protein modularity based on LWY repeat units. Neofunctionalization of LWY effector repertoire could be resulted from shuffling of functional modules. Effectors that adopt the PP2A-interacting module, formed by a specific (L)WY-LWY combination, gain the ability to competitively recruit the PP2A core enzyme in the host cells and facilitate infection. In these effectors, the PP2A-interacting module is combined with divergent LWY units in the C terminus, leading to functional diversity in the effector-PP2A holoenzymes. This model was created using Biorender.

### Limitations of the study

Our study demonstrates how LWY tandem repeat units in *Phytophthora* effectors can serve as functional modules and shows that one such module enables the effectors to hijack the protein phosphatase 2A (PP2A) core enzyme in the host. Effectors harbour the same PP2A-hijacking module have diverse LWY units in their C-terminus, indicating that domain shuffling events based on these tandem repeats may facilitate their functional diversification. Future studies are required to validate the contribution of C-terminal LWY units in these effectors to substrate specificity of the effector-PP2A holoenzymes. Furthermore, structural analysis of the holoenzymes, with information on substrate-binding site, will provide insight into the detailed interactions between effectors, substrates, and the PP2A catalytic subunit. Finally, the mechanism underlying the potential shuffling of these repeat units remains to be determined. The identification of PP2A core enzyme as a susceptibility factor that is targeted by viral, bacterial and oomycete pathogens warrant future research to precisely engineer this pathogen-target hub in order to disrupt pathogen virulence and generate resistance.

## ACKNOWLEDGMENTS

We thank the staff of the BL-17U1 and BL-19U1 beamlines at the Shanghai Synchrotron Radiation Facility, and the BL41XU beamline at SPring-8 (2019A2533). We thank staff at the John Innes Ccenter proteomics for help with implementation of FAIMS enabled LC-MS/MS. We thank Drs. Juan Dong and Cyril Zipfel for kindly providing Arabidopsis seeds of some of the PP2A mutants, Dr. Zhiyong Wang for providing the YFP-TurboID system, and Dr. Sophien Kamoun for the PITG_16705 construct. **Funding:** W. M. is supported by Gatsby Charitable Foundation, UKRI BBSRC Grant BBS/E/J/000PR9797, U.S. Department of Agriculture National Institute of Food and Agriculture grant 2018-67014-28488 (jointly offered by the National Science Foundation IOS-1758889). Y.W. is supported by grants from the Natural Science Foundation of China (31930065, 31725008, 22121003, 91940302, 32071444, 32071198 and 31630015), the Chinese Ministry of Science and Technology (2017YFA0504203), and the Chinese Academy of Sciences (XDB37010202).

## AUTHOR CONTRIBUTIONS

W.M. conceived the project. W.M. and Y.W. guided the execution of the experiments and oversaw the project. H.L., J.W., T.A.K., B.T., M.H., J.W., Z.C., Y.L., Y.Z., Y.H., J.S., P.D. and F.L.H.M. did the experiments and analyzed the data. H.L. and J.W. prepared figures and tables. W.M., H.L., J.W. and Y.W. wrote the manuscript with contributions from all authors.

## SUPPLEMEANTAL FIGURES

**Figure S1.**
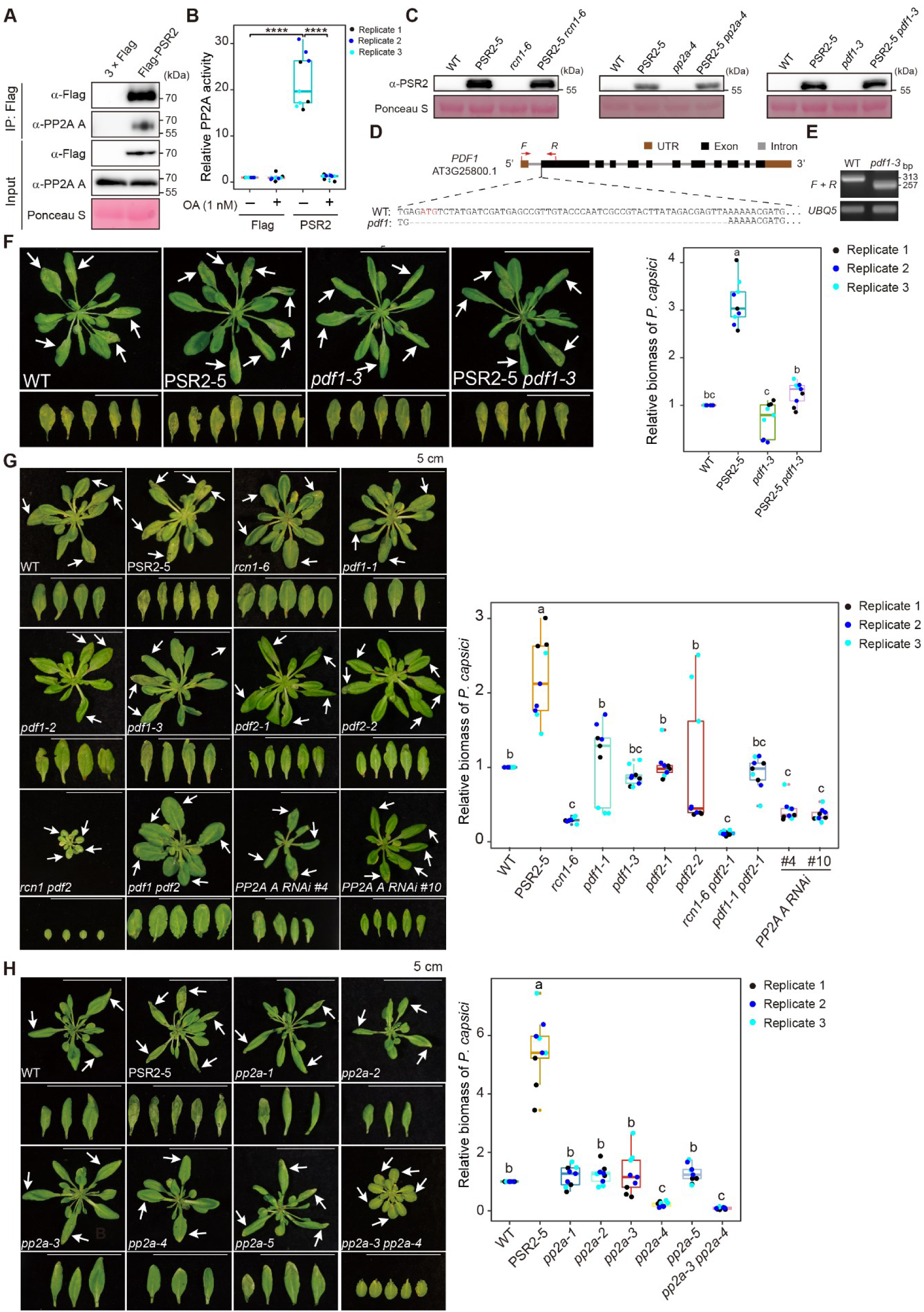
PSR2 forms a functional holoenzyme with PP2A core enzyme in *N. benthamiana* and this interaction is essential for the virulence activity of PSR2, related to Figure 1. (A) Co-immunoprecipitation of PSR2 with PP2A A subunit(s). *Agrobacterium*-mediated transient expression was used to express 3 × Flag-PSR2 in *N. benthamiana* leaves. Total proteins extracted from the infiltrated tissues were immunoprecipitated using anti-Flag magnetic beads. Co-precipitation of the PP2A A subunit(s) was detected using an anti-PP2A A antibody. Unlike in Arabidopsis where three bands representing three A subunit isoforms could be detected by the antibody, only one band was detectable from *N. benthamiana*, which may represent one major A subunit isoform or multiple isoforms with similar sizes. *N. benthamiana* infiltrated with *Agrobacterium* carrying the empty vector (3 × Flag) was used as a control. Equal loading was indicated by Ponceau S staining of the membrane. (B) PSR2 protein complex possessed PP2A phosphatase activity. Immunoprecipitated protein complexes from (A) were assayed for phosphatase activity, which was presented as fold changes compared to tissues infiltrated with *Agrobacterium* carrying the empty vector (3 × Flag). The phosphatase activity could be completely inhibited by Okadaic acid (OA), which specifically inhibits PP2A activity. Values from three independent experiments were analyzed by two-tailed Student t-test (\*\*\*\**p* < 0.0001). (C) Western blotting showing the protein levels of PSR2 in transgenic Arabidopsis plants. Total proteins were extracted from 2-week-old Arabidopsis seedlings and PSR2 was detected using an anti-PSR2 antibody. Ponceau S staining was used to confirm equal loading. (D) *pdf1* null mutant was generated in Arabidopsis by CRISPR/Cas9-based mutagenesis. The *pdf1-3* allele has a 56 nucleotides deletion in the first exon of the *PDF1* gene. (E) RT-PCR-based genotyping of the *pdf1-3* mutant allele using Ubiquitin 5 (*UBQ5*) as an internal control. Total RNAs were extracted from 14-day-old seedlings using TRIzol. Primes used to amplify the *PDF1* cDNA fragment were shown in (D). (F) Disease symptoms of Arabidopsis inoculated with *Phytophthora capsici* isolate LT263. Four-week-old plants were inoculated with zoospore suspensions of *P. capsici*. Photos were taken at 3 days post inoculation (dpi) with arrows indicating the inoculated leaves. Relative biomass of *P. capsici* was determined (n ≥ 20 in each sample per experiment) and data from three biological replicates are presented. One-way ANOVA and post hoc Tukey were used for statistical analysis. Different letters label significant differences (*p* < 0.05). WT, wild-type Col-0. Exact *P* values for all the experiments are provided in Supplementary Table 6. (G) Single and higher order mutants of the three PP2A A subunit genes (*RCN1*, *PDF1* and *PDF2*) exhibited either non-altered or enhanced resistance to *P. capsici*. The PP2A A RNAi line has all three A subunit genes knocked down. Knock out of all three genes results in lethality. The *rcn1 pdf1* double mutant has a severe developmental phenotype (data not shown) and could not be properly inoculated. Exact *P* values for all the experiments are provided in Table S6. (H) Single and higher order mutants of the five PP2A C subunit genes (*PP2A-1* to *PP2A-5*) exhibited either un-altered or enhanced resistance to *P. capsici*. Exact *P* values for all the experiments are provided in Table S6.

**Figure S2.**
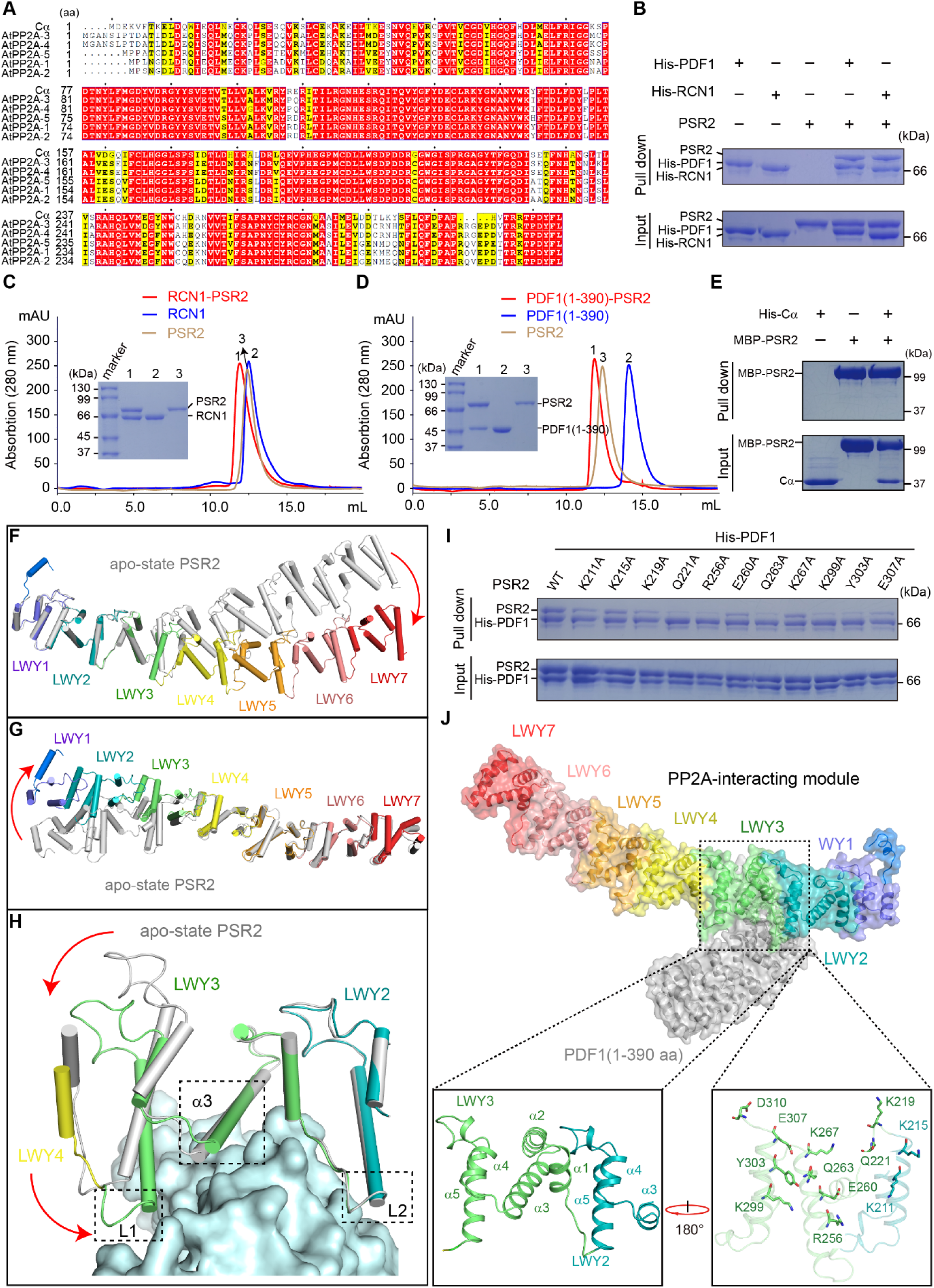
Structural and biochemical analysis of PSR2 interaction with PP2A A subunits, related to Figure 2. (A) Multiple sequence alignment showing sequence conservation between the human PP2A C subunit C**a** and the Arabidopsis PP2A C subunits. Because we could not obtain soluble proteins from any of the Arabidopsis PP2A C subunits, C**a** was used to test the direct interaction with PSR2. (B) PSR2 directly interacts with RCN1 and PDF1 *in vitro*. Protein composition was analysed by SDS-PAGE and Coomassie blue staining. (C) Gel filtration chromatography (in red) shows that RCN1 and PSR2 can form a binary complex. Chromatography of apo state PSR2 (in gold) and RCN1 (in blue) were used as controls. (D) Gel filtration chromatography (in red) shows the binary complex formed by PDF1(1-390 aa) and PSR2. Chromatography of apo state PSR2 and PDF1 (1-390 aa) were used as controls. (E) *In vitro* pull-down assay shows that PSR2 does not directly interact with C**a**. (F) Structural superimposition between binary- and apo state PSR2 based on the alpha carbon of amino acids range from 61-217 aa shows the conformational change triggered by PSR2-PDF1 interaction. (G) Structural superimposition between binary- and apo state PSR2 based on the alpha carbon of amino acids range from 311-670 aa shows the conformational change triggered by PSR2-PDF1 interaction. (H) Structural superimposition between binary and apo state PP2A-interacting module based on the alpha carbon of amino acids range from 180-250 aa shows PP2A-interacting module in PSR2 undergoes a significant conformational change with the L1 moving toward PDF1, thus holding PDF1 tightly. (I) Results of *in vitro* pull-down assays used to examine individual mutants, each has one of the 11 residues replaced with an alanine, for their contribution to interaction with PDF1. (J) Characterization of the PP2A-interacting module on the PSR2 identified two residues in LWY2 (in teal, K211 and K215 in α5) and ten in LWY3 (in lime, K219, Q221, R256, E260, Q263, K267, K299, Y303, E307, and D310 spread in all five α-helices) that directly mediate interaction with PDF1.

**Figure S3.**
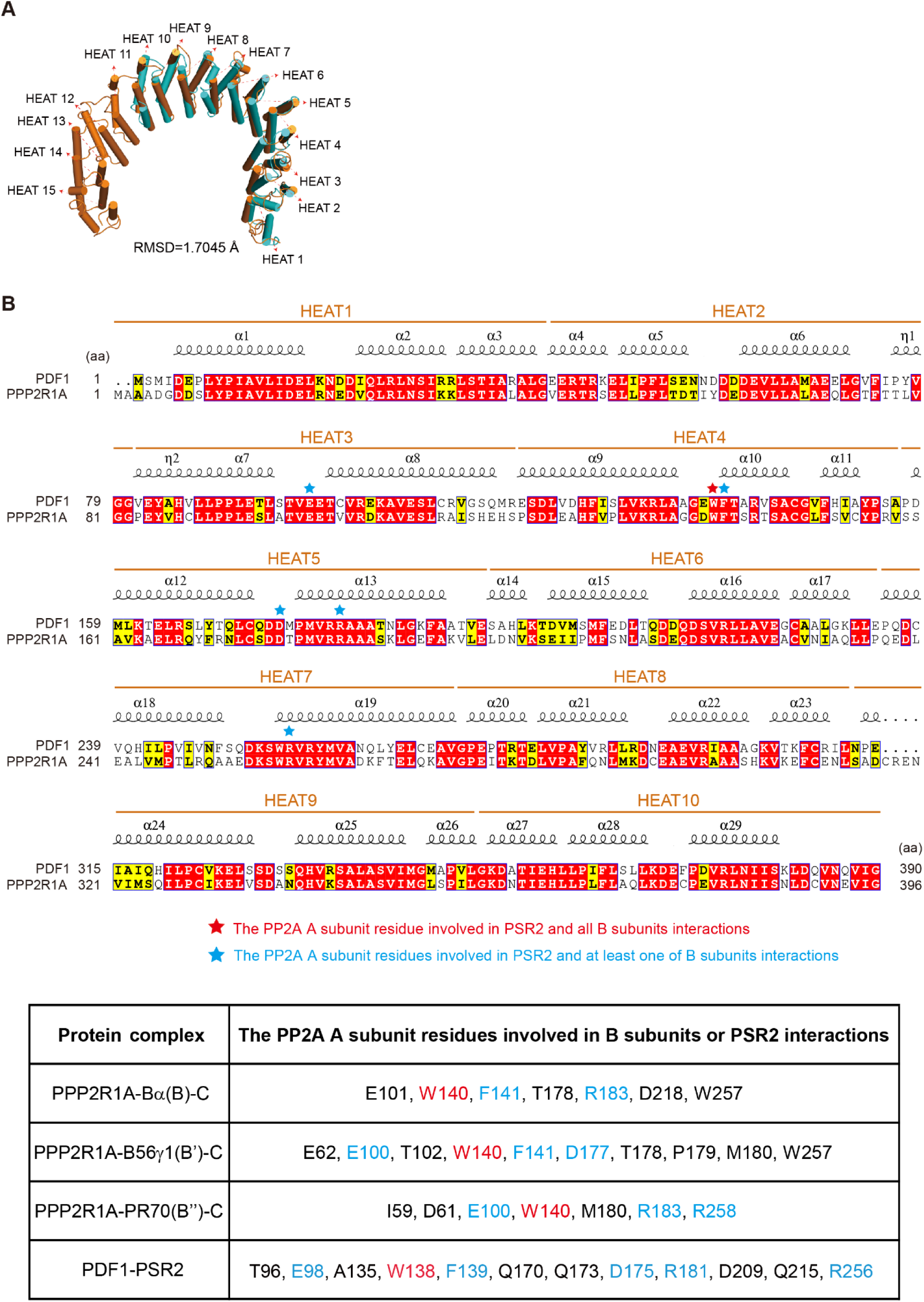
Similar residues in PDF1 and the human PP2A A subunit PPP2R1A are involved in interactions with PSR2 and human B subunits respectively, related to Figure 3. (A) Structural superimposition of PDF1 (in cyan) from the PDF1(1-390 aa)-PSR2(59-670 aa) complex and the A subunit PPP2R1A (in orange) from the human PP2A holoenzyme (PDB code:2iae) shows a high level of structural similarity. (B) Sequence alignment of PDF1 (1-390 aa) and PPP2R1A (1-396 aa) with residues involved in the interaction with PSR2 and endogenous human B subunits highlighted by asterisks. The Tryptophan residue (in red) is involved in all interactions and the residues in blue are involved in PDF1-PSR2 interaction as well as at least one of the three interactions in the human PP2A holoenzymes. The table shows the PP2A A subunit residues involved in interactions with B subunits or PSR2 in four protein complexes.

**Figure S4.**
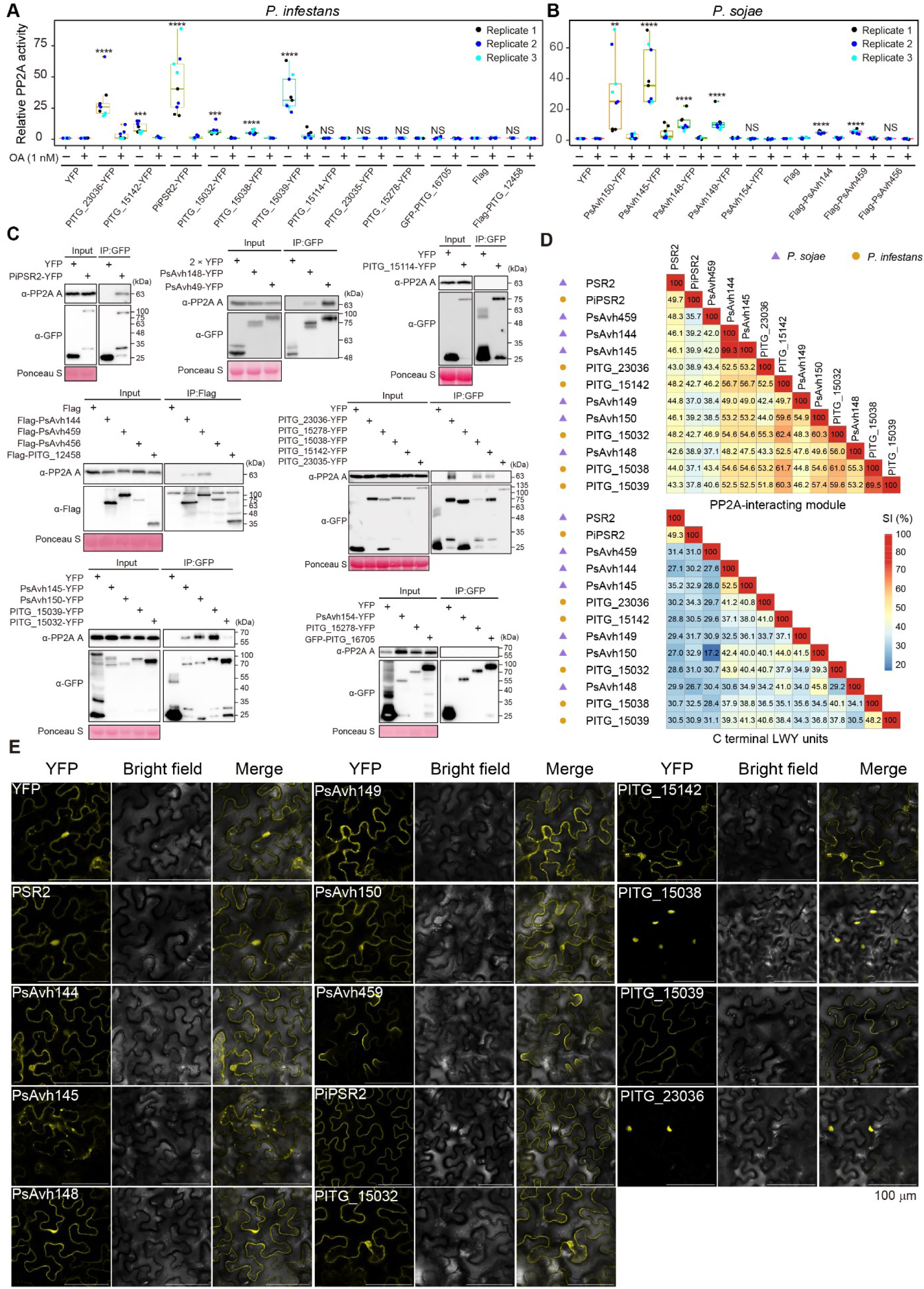
Twelve LWY effectors, in addition to PSR2, can associate with functional PP2A core enzyme in *N. benthamiana*, but exhibit diversification in their C-terminal LWY units related to Figure 4. (A and B) Phosphatase activity was determined from complexes formed by *P. infestans* effectors (A) or *P. sojae* effectors (B). Individual effectors were transiently expressed in *N. benthamiana*. Effector complexes were immunoprecipitated using anti-GFP or anti-Flag magnetic beads and subsequently subjected to measurement of phosphatase activity. Okadaic acid (OA) specifically inhibits PP2A phosphatase activity. Values are from three biological replicates and analzyed by two-tailed Student t-test. Asterisks label values with statistically significant differences (\*\*\**p* < 0.001, \*\*\*\**p* < 0.0001). Exact *P* values for all the experiments are provided in Table S6. (C) PP2A A subunit(s) was co-immunoprecipitated with the effectors that could form complexes with PP2A phosphatase activity. Western blotting using the co-immunoprecipitated samples in (A and B) was used to detect PP2A A subunit(s) by an anti-PP2A A antibody. *N. benthamiana* tissues expressing 2 × YFP or infiltrated with *Agrobacterium* carrying the empty vector (3 × Flag, YFP or GFP) were used as controls. Equal loading of the samples was confirmed by Ponceau S staining of the membranes. (D) C-terminal LWY units exhibit a higher level of diversity in the PP2A-associating effectors. Sequence identities are represented by percentage amino acid identities calculated by pairwise sequence alignment (EMBOSS needle). (E) Subcellular localization of PP2A-interacting effectors when transiently expressed in *N. benthamiana*. The effectors are C-terminally tagged with yellow fluorescent protein (YFP). Bars, 100 **m**m.

**Figure S5.**
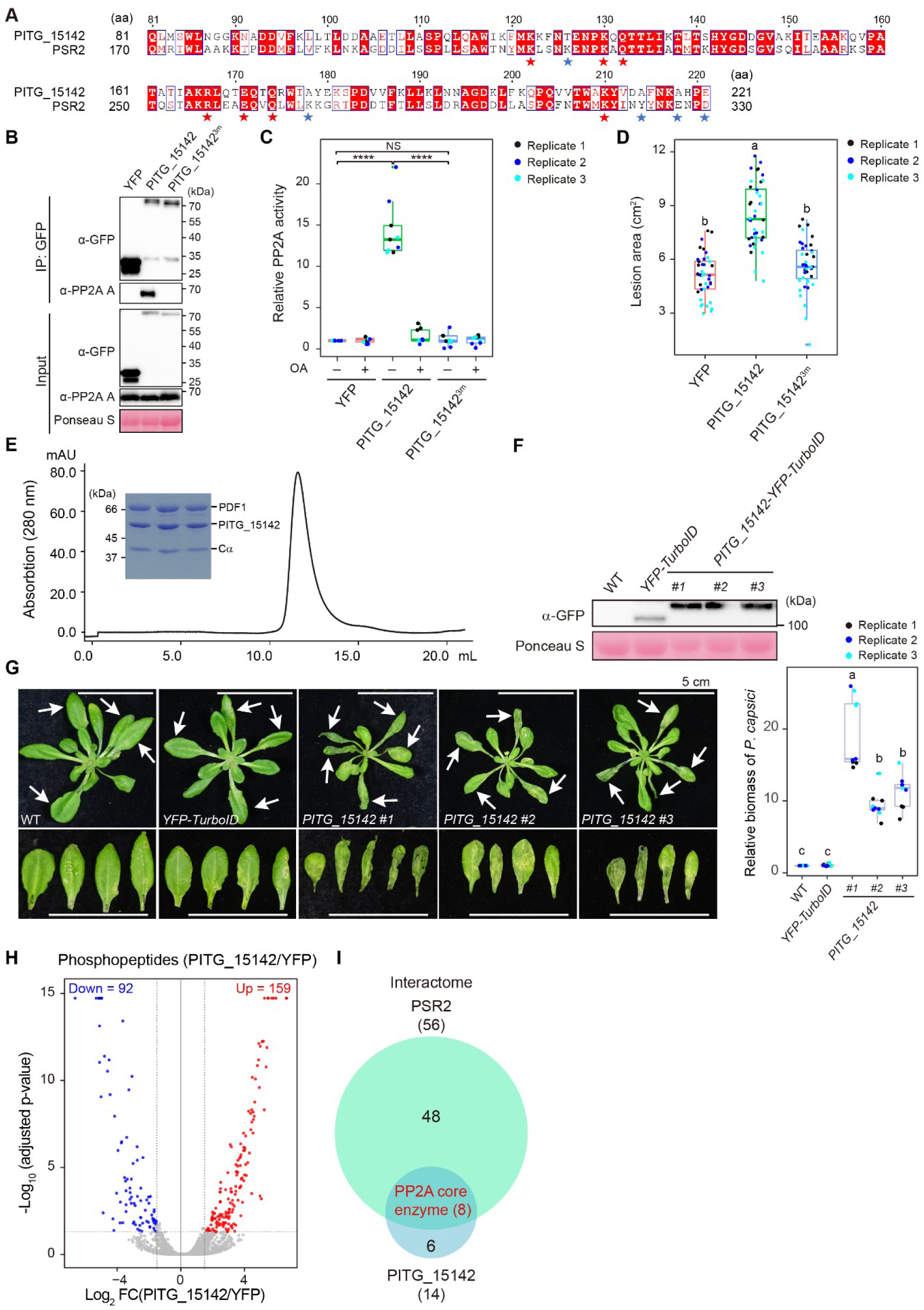
PITG_15142 hijacks the host PP2A core enzyme through a conserved mechanism with PSR2 but regulates a distinct set of host proteins, related to Figure 5. (A) Sequence alignment between the PP2A-interacting module in PITG_15142(81-221 aa) and PSR2(170-310 aa). Asterisks label the 12 residues of PSR2 that have direct contacts with PDF1. Red asterisks highlight the conserved residues in PITG_15142. (B and C) The REQ triad is required for the formation of a functional PITG_15142-PP2A holoenzyme *in planta*. YFP-tagged PITG_15142 or PITG_15142^R167A/E171A/Q174A^ (PITG_15142^3m^ in the figure) were expressed in *Nicotiana benthamiana* and immunoprecipitated using anti-GFP magnetic beads. Co-immunoprecipitation of PP2A A subunit(s) was detected using an anti-PP2A A antibody (B) and the phosphatase activity was measured with or without the PP2A inhibitor Okadaic acid (OA) (C). For the phosphatase assay, values obtained in three independent biological repeats were analyzed by two-tailed Student t-test (\*\*\*\**p* < 0.0001; NS, not significant.). *P* values for all the experiments are provided in Table S6. (D) The REQ triad is required for the virulence activity of PITG_15142. YFP-tagged PITG_15142 or PITG_15142^3m^ were expressed in *Nicotiana benthamiana*. 48 hours after Agro-infiltration, the leaves were inoculated with mycelium plugs of *P. capsici*. Lesion areas were quantified using Image J program at 3 days post inoculation. Data from three biological replicates are presented (n `≥` 10 in each sample per experiment). One-way ANOVA and post hoc Tukey were used for statistical analysis. Different letters label significant differences (*p* < 0.05). Exact *P* values for all the experiments are provided in Supplementary Table 6. (E) Gel filtration chromatography shows the formation of PDF1-PITG_15142-Cα complex *in vitro*. (F) Western blotting confirming YFP or PITG_15142 protein expression in the transgenic Arabidopsis. All constructs have the C-terminal YFP-TurboID tag. Total proteins were extracted from 2-week-old seedlings and examined using an anti-GFP antibody. Ponceau S staining was used to assess equal loading. (G) Expression of PITG_15142 increased the susceptibility of Arabidopsis to *P. capsici*. Four-week-old plants were inoculated with zoospore suspensions. Photos were taken at 3 days post inoculation with arrows indicating inoculated leaves. Relative biomass of *P. capsici* was determined (n ≥ 20 in each sample per experiment) and data from three biological replicates are presented. One-way ANOVA and post hoc Tukey were used for statistical analysis. Different letters label significant differences (*p* < 0.05). WT, wild-type Col-0. Exact *P* values for all the experiments are provided in Supplementary Table 6. (H) Volcano plots show changes in the phosphopeptide abundance according to the average ratio (log_2_) and *p* value (-log_10_ adjusted *p*-value) in Arabidopsis expressing PITG_15142-YFP-TurboID. Gray dots represent phosphopeptides with non-significantly change in abundance. Red and blue dots represent phosphopeptides with significantly increased and decreased abundance, respectively. The vertical dashed lines indicate *p*-value = 0.05 and the horizontal dashed line indicates a fold change of 2. (I) A comparison of the PSR2 and PITG_15142 interactomes analyzed by IP-MS in Arabidopsis. The only proteins that commonly interacted with both effectors were the three PP2A A subunits and the five C subunits.

## STAR METHODS

### CONTACT FOR REAGENT AND RESOURCE SHARING

Further information and requests for resources and reagents should be directed to and will be fulfilled by the Lead Contact, Wenbo Ma or Yanli Wang

### DATA AND MATERIAL AVAILABILITY

The crystal structure data of PDF1-PSR2 and PITG_15142 have been deposited in https://deposit-pdbj.wwpdb.org/deposition. The mass spectrometry proteomics data have been deposited to the ProteomeXchange Consortium via the PRIDE (Perez-Riverol et al., 2019) partner repository with dataset identifiers PXDxxxxx and xxxxxxxxxxx. Materials are available from W.M. upon request under a materials transfer agreement with the Sainsbury Laboratory.

### METHOD DETAILS

#### Plant materials and growth conditions

*Arabidopsis thaliana* and *Nicotiana benthamiana* plants were grown in a growth room at 22°C with a 16/8h light/dark regime. *Arabidopsis thaliana* accession Col-0 was used as wild-type and for generating transgenic plants. Sterile Arabidopsis seedlings were grown on plates containing Murashige-Skoog medium and 1% sucrose supplemented with 0.8% Phytagel in a growth chamber with the setting of 22°C and a 16/8 h light/dark regime. Arabidopsis mutants and transgenic plants (Bian et al., 2020; Hsu et al., 2019; Qiao et al., 2013; Segonzac et al., 2014) used in this study are listed in the key resources table. Primers used to genotype the *pp2a* mutants are listed in the key resources table.

#### Plasmid construction

All the LWY effectors were cloned without their N-terminal secretion signal peptide for various experiments. Using the LR Clonase II-based gateway cloning system, the effector genes were first cloned into the pENTR/D-TOPO vector (Invitrogen) and then destination vectors (pEarleyGate100 or pEarleyGate101, Invitrogen) for *in planta* expression (Key resources table). Some effectors were also cloned into the pGW514 vector by In-fushion system for *in planta* expression. For expression in *E. coli*, effectors or PP2A subunits were cloned into vectors that add N-terminal 6×His tags to facilitate protein purification using Ni-NTA Sepharose resin. Primers used to amplify these sequences are listed in key resources table.

#### Generation of Arabidopsis mutants and transgenic plants

To generate a *pdf1* knockout mutant in Arabidopsis, CRISPR/Cas9-based mutagenesis was used. A guide RNA was designed to target to the first exon of the *PDF1* gene using the Optimized CRISPR Design-MIT website (http://crispr.mit.edu/). This guide RNA was introduced into the *YAO* Promoter-Driven CRISPR/Cas9 vector and delivered into *Agrobacterium tumefaciens* strain GV3101 for Arabidopsis transformation (Clough and Bent, 1998; Yan et al., 2015). The resulting transgenic T1 seeds were screened on ½MS medium with hygromycin (Roche). Genomic fragments covering the target site were sequenced to confirm mutation.

To obtain PITG_15142-YFP-TurboID transgenic plants, *A. tumefaciens* carrying the construct p35S::PITG_15142-YFP-TurboID was used for Arabidopsis transformation. The resulting transgenic T1 seeds were screened on ½MS medium with Phosphinothricin (Duchefa).

#### Antibody production

Full length RCN1 proteins with a Glutathione S-transferase (GST) tag were purified after expression in *E. coli* used to generate polyclonal antibodies in rabbits (produced and purified by Pacific Immunology Corp). The antibody is able to detect all three PP2A A subunits of Arabidopsis in distinctive bands through western blotting but only one band using leaf tissues of *N. benthamiana*.

#### Immunoprecipitation of effector protein complexes

Two grams of leaf tissues from two-week-old wild-type or PSR2-expressing Arabidopsis (Hou et al., 2019; Qiao et al., 2013) seedlings were ground in liquid nitrogen and suspended in 2 mL IP buffer (10% (v/v) Glycerol, 50 mM Tris-HCl pH 7.5, 50 mM NaCl, 1 mM EDTA, 1×protease inhibitor mixture, 5 mM DTT, 2% PVPP and 0.1% NP-40). Samples were centrifuged at 13000 rpm for 15 min at 4°C. Supernatants were incubated with 10 **m**g of an anti-PSR2 antibody (Hou et al., 2019) for 2 hr at 4°C with agitation. 25 **m**L of pre-washed Protein A magnetic beads (Thermo Scientific) were added into the mixture and incubated for another 2 hr at 4°C. Beads were washed three times (10% (v/v) Glycerol, 50 mM Tris-HCl pH 7.5, 50 mM NaCl, 1 mM EDTA, and 0.1% NP-40). Bound PSR2 and its associated proteins were separated by SDS-PAGE electrophoresis for western blotting or Mass Spectrometry (MS) analysis. In western blotting, PSR2 or PP2A A subunits were detected using anti-PSR2 and anti-PP2A A antibodies respectively. Details in MS analysis are described in a separate section below.

The interaction proteins of PITG_15142 were identified using the same method using leaf tissue from transgenic Arabidopsis seedlings expressing PITG-15142-YFP-TurboID. Leaf tissues from transgenic plants expressing YFP-TurboID (Hsu et al., 2019) were as the control. GFP-Trap magnetic agarose (Chromotek) was used to enrich the effector protein complexes.

Interaction of LWY effectors with PP2A A subunits were also detected in *N. benthamiana* using a similar procedure. *A. tumefaciens* carrying constructs for expressing the LWY effectors was infiltrated into leaves of 3-week-old plants. The effectors were immune-precipitated with anti-GFP (Clontech) or anti-Flag antibody (Sigma-Aldrich). PP2A A subunits were detected using the anti-PP2A A antibody by western blotting.

#### Mass spectrometry and data analysis

Immunoprecipitated samples were separated on SDS-PAGE and stained with colloid Coomassie Brilliant Blue (Simple stain, Invitrogen). Gel slices were cut into small pieces and de-stained in 20% Acetonitrile by repeated washing. Cysteine residues were modified by 30 min reduction in 10 mM DTT followed by 20 min alkylation using 50 mM chloroacetamide. Then extensive washing and dehydration were performed with 20% and 100% Acetonitrile respectively. The gel slices were incubated with 100 ng of trypsin (Promega) in 50 mM ammonium bicarbonate, and 10% Acetonitrile at 37°C overnight. The digestion was stopped with an equal volume of 1% formic acid in 25% Acetonitrile. Peptides were extracted three times using 25% Acetonitrile, evaporated to dryness in a rotary vacuum evaporator, and stored at −20°C.

LC-MS/MS analysis was performed using a hybrid mass spectrometer Orbitrap Fusion and a nanoflow UHPLC system U3000 (Thermo Scientific). Tryptic peptides, dissolved in 2% Acetonitrile, and 0.2% Trifluoroacetic acid (TFA), were injected onto a reverse-phase trap column nanoEase M/Z Symmetry C18, beads diameter 5 µm, 180 **m**m × 20mm (Waters, Corp.). The column was operated at the flow rate of 20 **m**l/min in 2% Acetonitrile, 0.05% TFA. After 2.5 min the trap column was connected to the analytical column nanoEase M/Z HSS C18 T3 Column, beads diameter 1.8 µm, 75 **m**m × 250mm (Waters). The column equilibrated with 3% B buffer before the injection in 3% B (B buffer: 80% Acetonitrile in 0.05% FA) was subsequently eluted with the linear-gradient of B buffer. The flow rate was set to 200 nL/min. The mass spectrometer was operated in positive ion mode with a nano-electrospray ion source. Molecular ions were generated by applying voltage +2.2 kV to a conductive union coupling the column outlet with fussed silica PicoTip emitter, ID 10µm (New Objective, Inc.). The ion transfer capillary temperature was set to 275°C and the focusing voltages in the ion optics were in the factory default setting.

A method for mass spectrometer has been designed and tested with maximum sensitivity gain for samples of low complexity, such as immunoaffinity enriched protein complexes from plants. MS events consisted of a full scan in an Orbitrap analyser followed by two collisions of “soft” CID (collision-induced dissociation) and more “energetic” HCD (Higher-energy collisional dissociation) to maximize the chances to acquire spectra with structurally important information. The fragment ions were detected with the low-resolution detector at the ion trap end.

Fusion Software v3.3 was installed. Orbitrap full scan resolution was set to 120,000, mass range m/z 300 to 1800 automatic gain control (AGC) for the target 200,000 ions and maximal infusion time 50 ms. The precursor dissociation events were driven by a “data-dependent algorithm” (DDA) with the dynamic exclusion 20 s after the collision had been triggered. The number of MS/MS events was the maximum possible between full scans with the frequency of not more than 3 s period (“top speed” settings). Gain settings were AGC = 10,000 for the Ion Trap, maximal injection time = 35 ms. The isolation width and normalized collision energy for both collision events CID and HCD were set to m/z 1.6 and CE = 30% respectively. Only precursor ions with a positive charge state 2-7 and an intensity threshold greater than 10,000 were submitted to fragmentation.

Peak lists in the format of Mascot generic files (mgf files) were generated from raw data files using MS Convert (ProteoWizard 3.0.9740) and sent to a peptide search on Mascot server v.2.7 using Mascot Daemon (Matrix Science, Ltd.). The lists were searched against protein databases including typical proteomics contaminants such as keratins, etc. Tryptic peptides with up to 2 possible mis-cleavages and charge states +2, +3, and +4 were allowed in the search. The following peptide modifications were included in the search: carbamidomethylated Cysteine (static) and oxidized Methionine (variable). Data were searched with a monoisotopic precursor and fragment ion mass tolerance of 10 ppm and 0.6 Da respectively. The decoy database was used to validate peptide sequence matches. Mascot results were combined in Scaffold v5.1.0 (Proteome Software Inc.) and exported to Excel (Microsoft) for further processing and comparisons(Searle, 2010).

The probability filter in Scaffold was set to 1% FDR (false discovery rate) for both peptide and protein identifications, with at least 2 unique peptides identified per protein. Protein probabilities were calculated in Scaffold by the Protein Prophet algorithm; proteins that contained similar peptides and could not be differentiated based on MS/MS analysis alone were grouped to satisfy the principles of parsimony. Fold change analysis was carried out followingly. Total spectral counts (SPC) values of all the sample types and replicates were exported from the Scaffold for proteins and peptides above 1% FDR. The contaminants removed; (keratin, trypsin, BSA, etc.) From binary comparisons were removed hits that showed missing values in our target protein measurements (the proteins baits in our immunoaffinity preparations). Any missing values and zeros (low scoring hits) were replaced by an arbitrary number of 0.001 to allow for the ratio calculation and visualisation purposes. Log_2_ ratios of averaged SPC in selected binary comparisons were calculated. Data were sorted and plotted in a bar chart. The enrichment ratio two-fold or better was considered as significant.

#### PP2A phosphatase activity assay

For measuring the phosphatase activity of PSR2 complex in *Arabidopsis*, proteins were enriched from leaf tissues using the anti-PSR2 antibody as described above. PP2A phosphatase activity was measured using a non-radioactive molybdate dye-based phosphatase assay kit (#V2460, Promega) in which a synthetic phosphopeptide (RRA[pT]VA) was used as the substrate. A reaction mixture containing 100 **m**M phosphopeptide and immunoprecipitated proteins was incubated at 37°C for 5 min with or without 1 nM Okadaic acid, which is a PP2A-specific inhibitor(Bialojan and Takai, 1988). An equal volume of molybdate dye-additive was used to stop the reaction. Phosphate released from the phosphopeptide was measured as absorbance at 600 nm against a standard curve. Relative PP2A activity was calculated as the ratio between the experimental group and the control (WT plant or plants harboring the empty vector).

For measuring the phosphatase activity of effector complexes in *N. benthamiana*, effector proteins were transiently expressed in leaves using Agro-infiltration and enriched from leaf tissues using GFP-Trap magnetic agarose or anti-Flag magnetic beads. Relative PP2A activity was examined as described above.

#### Arabidopsis and *N. benthamiana* infection assays by *Phytophthora capsici*

Inoculation using *P. capsici* isolate LT263 was performed as previously described (Hou et al., 2019; Wang et al., 2013). Arabidopsis leaves were inoculated with zoospore suspension and *N. benthamiana* leaves were inoculated with mycelium plugs.

For Arabidopsis inoculation, LT263 was grown on 10% V8 medium at 25°C in the dark until mycelia covered the whole plate. To induce sporulation, mycelium plugs were first washed and then incubated with sterile tap water at 25°C for 24 hours in the dark. Zoospore release was induced by 4°C incubation for 40 min, followed by light induction for 20 min at room temperature. Zoospores were collected using one layer of miracloth (Millipore) for making suspensions (200–500 zoospores/µL) to be used for inoculation. 3-6 adult rosette leaves per Arabidopsis plant were inoculated using ∼20 µL zoospore suspension applied to the abaxial side of each leaf. Leaves treated with water were used as a mock control. The inoculated plants were placed in a growth chamber with a transparent cover to keep high humidity. Disease symptoms were monitored three days after inoculation and DNA was extracted from all the inoculated leaves (n ≥ 20 per treatment). The biomass of *P. capsici* was determined by qPCR using *P. capsici* specific primers (listed in the key resources table). The Arabidopsis *rub4* was used an internal control. Relative biomass of *P. capsici* in mutant or transgenic plants was determined by comparing to the value from the biomass in wildtype (WT) plants, which was set as “1”.

For *N. benthamiana* inoculation, the abaxial sides of detached leaves were inoculated with fresh mycelial plugs (0.5 cm). Leaves were kept in sealed 0.8% water agar plates in the dark at 25°C. Lesions were observed under UV light three days after inoculation. Sizes of the lesion areas were analyzed using imageJ (https://imagej.net/).

#### Protein expression and purification for structural analysis

All proteins, mutants and truncations were expressed in *E. coli* BL21(DE3) (Novagen). Transformants carrying the recombinant plasmids were induced by 0.1 mM IPTG at OD_600_ = 0.6 for 16 hr at 16°C. Cells were lysed by sonication in a lysis buffer (20 mM Tris-HCl pH 7.5, 0.5 M NaCl) at 4°C, and target proteins were purified by a Ni-NTA Sepharose resin column (GE Healthcare) and further fractionated by ion exchange (GE Healthcare). Proteins with the 6 × His-Sumo tag were digested with ubiquitin-like protein 1 (Ulp1) protease and dialyzed against a buffer containing 20 mM Tris-HCl (pH 7.5), 0.3 M NaCl for 2 hr at 4°C. The proteins were further purified by the Ni-NTA Sepharose resin column to remove the 6 × His-Sumo tag. The flow-throughs were collected for fractionation by ion exchange.

We tried to express three Arabidopsis PP2A B subunits, ATB**b**, ATB’**a**, and ATB’**g,** in *E. coli*. ATB**b** could not be expressed, ATB’**a** proteins were unstable during protein purification. Only ATB’**g** can be purified and used for further experimentation.

None of the Arabidopsis PP2A C subunits could be expressed despite numerous trials using various expression systems. Therefore, we worked with a human PP2A C subunit (C**a**) and obtained purified proteins after expression in baculovirus-infected Hi-5 suspension culture as described previously (Cho and Xu, 2007).

#### Crystallization, data collection and structure determination

PDF1-PSR2 and RCN1-PSR2 were used for extensive screening of crystallization conditions. The hanging-drop vapor-diffusion method was used for crystal growth. All crystals were obtained by mixing 1 **m**L PDF1-PSR2 complex with 1 **m**L reservoir solution and incubated at 16°C. Using full-length PDF1, we only determined the crystal structure of its binary complex with PSR2 at a resolution of 4.2 Å. To improve the resolution, we used truncated forms of PDF1 and PSR2, and finally acquired high resolution crystal structure of the PSR2(59-670 aa)-PDF1(1-390 aa) complex for structural analysis. The crystals of this complex were grown using the condition of 0.1 M Tris-Bis pH 5.8, 20 mM MgSO_4_, and 5% PEG 20000. The crystals of PITG_15142 were grown from 0.1 M Tris-Bis propane pH 7.5, 1 mM CdCl_2_ and 19.5 % PEG 600. All crystals were flash-frozen in liquid nitrogen with 20% glycerol. Diffraction datasets were collected at beamline BL17U1 or BL19U1 at Shanghai Synchrotron Radiation Facility (SSRF), or at beamline BL41XU at Spring-8 in Japan, and processed with XDS or HKL2000 (Otwinowski and Minor, 1997). All structures were solved by molecular replacement (MR) using PHENIX PHASER(McCoy et al., 2007) and built manually in COOT (Emsley et al., 2010). The search model for the LWY effectors was a predicted structure by AlphaFOLD2(Jumper *et al*., 2021). Iterative cycles of crystallographic refinement were performed using PHENIX (Adams et al., 2002). All structural figures were prepared using PyMOL (http://www.pymol.org/).

#### Reconstitution of PDF1-PSR2, RCN1-PSR2, PDF1-Ca-PSR2, PDF1-Ca-ATB’g and PDF1-Ca-PITG_15142 protein complexes

To get the PDF1-PSR2 or RCN1-PSR2 binary complex, PDF1/RCN1 and PSR2 were mixed in the reaction buffer (20 mM Tris-HCl pH 7.5, 200 mM NaCl 1 mM DTT) at a molar ratio of 1:1, incubated on ice for 15 min, and purified by gel filtration chromatography (Superdex 200 increase 10/300GL, GE Healthcare). For the PDF1-C**a**-PSR2, PDF1-C**a**-ATB’**g** or PDF1-C**a**-PITG_15142 complexes, PDF1, C**a** and PSR2/ATB’**g**/PITG_15142 were mixed in reaction buffer (20 mM Tris-HCl pH 7.5, 1mM DTT) with 300/300/150 mM NaCl, respectively, at a molar ratio of 1: 1: 1. Purified proteins were added on at the time in the order listed and incubated for 15 min on ice between proteins. The complexes were then purified by gel filtration chromatography and analyzed using SDS-PAGE.

#### Isothermal titration calorimetry (ITC)

ITC was performed using the ITC200 microcalorimeter (MicroCal, GE Healthcare) at 16°C by setting 600 rpm/min. The purified PDF1, ATB’**g** and PSR2 proteins were dialyzed against the ITC buffer (25mM HEPES pH 7.5, 200 mM NaCl, 5% glycerol) at 4°C. PSR2 or ATB’**g** was diluted to 16.5 **m**M, and then loaded into the cell chamber of the microcalorimeter. PDF1 were diluted to 165 **m**M and then loaded into the springe. Fourteen injections of PDF1 were carried out with a spacing of 120 s. At least two independent experiments were performed. The data were analyzed using Origin 9.0 (http://www.originlab.com/). Minor protein precipitation happened in the cell chamber because of rotation of the springe that may affect the K_d_ values, especially that of ATB’**g**.

#### *In vitro* Pull-down assay

6×His-PDF1 proteins were incubated with untagged wild-type or mutant PSR2 at a molar ratio of 1:1 in the buffer (20 mM Tris-HCl pH 7.5, 100 mM NaCl) on ice for 30 min. PDF1 protein complexes were pulled down using Ni-NTA Sepharose resins with non-specifically bound proteins removed by a wash buffer (20 mM Tris-HCl pH 7.5, 100 mM NaCl, 40 mM imidazole). The bound proteins were eluted by the elution buffer (20 mM Tris-HCl pH 7.5, 100 mM NaCl, 500 mM imidazole) and flow-through collections were analyzed by SDS-PAGE.

To investigate whether PSR2 can directly interact with PP2A C subunit(s), MBP-PSR2 were incubated with C**a** at a molar ratio of 1:1 in the buffer (20 mM Tris-HCl pH 7.5, 100 mM NaCl) on ice for 30 min. PSR2 was precipitated using Amylose resin (NEW ENGLAND BioLabs) and non-specifically bound proteins were removed by the wash buffer (20 mM Tris-HCl pH 7.5, 100 mM NaCl). The bound proteins were eluted by the elution buffer (20 mM Tris-HCl pH 7.5, 100 mM NaCl, 20 mM maltose) and flow-through collections were analyzed by SDS-PAGE.

#### Competition of PSR2 with ATB’g on recruiting PDF1-Cα core enzyme

A Superdex 200 Increase 10/300 gel filtration column (GE Healthcare) was used at a flow rate of 0.2 mL/min and with the absorbance monitored at 280 nm. Preformed PDF1-Cα-ATB’**g** complexes were prepared through incubating purified PDF1, C**a** and ATB’**g** at a molar ratio of 1: 1: 1. Components were added in the order listed and incubated for 15 min on ice before adding the next component. Then, equimolar PSR2 proteins were added to sample and incubated for another 15 min on ice. 100 **m**L of the sample was applied to pre-equilibrated column (20 mM Tris-HCl pH 7.5, 300 mM NaCl, 1 mM DTT) and analyzed by gel filtration chromatography. Fractions were also analyzed on a 12% SDS-PAGE gel and visualized by Coomassie blue staining. We also tested whether ATB’**g** could facilitate the dissociation of PSR2 from a preformed PDF1-C**a**-PSR2 complex using a similar procedure.

#### Identification of additional LWY effectors with the PSR2-like PP2A interacting module

Structural prediction of 80 LWY effectors from *P. infestans* and 54 LWY effectors from *P. sojae* (He et al., 2019) was conducted by AlphaFOLD2 (Jumper et al., 2021). The architecture of effectors in Figure 4A was created by Biorender (http://biorender.com) Next, sequences and structures of all the predicted (L)WY-LWY motifs were extracted using PyMOL (http://www.pymol.org/). Similarity analysis with the (L)WY2-LWY3 of PSR2 was performed by sequence-based analysis using EMBOSS Needle (pairwise sequence alignment, https://www.ebi.ac.uk/Tools/psa/emboss_needle/) and structure-based analysis. Structural superposition between the WY-LWY motifs with PP2A-interacting module of PSR2 was generated by aligning PDB files using fr-tm-align (Pandit and Skolnick, 2008). The corresponding residues at the positions of the 12 residues in PSR2 that directly mediate the interaction with PDF1 were pulled out from all the WY-LWY motifs. 12 PDF1-interacting residues of PSR2 from the alignment, TM-score, and root-mean-square deviation (RMSD) scores were extracted using bash scripting. Candidates from these analyses were further investigated for conservation on these residues. A conservation score was assigned to each residue using BLOSUM 62 and the sum value of all 12 scores was used to evaluate overall conservation of the candidate with (L)WY2-LWY3 of PSR2.

#### Phylogeny, structure and sequence analysis of PDF1-interacting module and the C terminal units of PP2A-interacting effectors

The amino acid sequences of the PDF1-interacting modules were aligned using MUSCLE in MEGA10. An unrooted phylogenetic tree was generated based on the aligned datasets using a maximum-likelihood method with the bootstrap, uniform rates, and use all sites in MEGA10. The tree is drawn to scale, with branch lengths measured in the number of substitutions per site. Sequence and structural similarity analysis was conducted using PDB file alignment by fr-tm-align and EMBOSS Needle, respectively as described above.

#### Subcellular localization of PP2A-interacting effectors

*Agrobacterium tumefaciens* carrying YFP or individual PP2A-interacting effectors tagged with C-terminal YFP constructs was infiltrated into leaves of *N. benthamiana* together with another *Agrobacterium* strain carrying the viral RNA silencing suppressor P19. All genes were under the control of CaMV *35S* promoter. At 3 days after inoculation, yellow fluorescence in the infiltrated leaves was observed under a confocal microscope (Leica SP8) (excitation at 488 nm and emission at 500-550 nm).

#### Phosphoproteomic analysis

Phosphopeptide samples were analysed using an Orbitrap Eclipse™ Tribrid™ Mass Spectrometer (Thermo Fisher Scientific) fitted with High-Field Asymmetric Waveform Ion Mobility Spectrometry (FAIMS) Pro Duo Interface and coupled to a U3000 nano-UPLC (Thermo Fisher Scientific). The dissolved peptides were injected onto a reverse phase trap column PepMap™ Neo Trap Cartridge (Thermo Scientific). Trap column flowrate was 20 μl/min in 2% acetonitrile, 0.05% TFA. Peptides were eluted from trap column onto the analytical column nanoEase M/Z HSS T3 Column, beads diameter 1.8 μm, inner diameter 75 μm x 250 mm length (Waters). The column was equilibrated with 3% B (B: 80% acetonitrile in 0.05% formic acid (FA), A: 0.1% FA) before subsequent elution with sigmodal gradient to 32% B over 100 min followed by 48 min gradient to 50% B and 10 min 99% B, for a total of 162 min. The flow rate was set to 200 nl/min. The mass spectrometer was operated in positive ion mode with nano-electrospray ion source with FAIMS enabled and scanning at three different CV values (−35, −50 and −65 v) per scan cycle (1 sec per CV for total of 3 sec). Molecular ions were generated by applying voltage +2.8kV to a conductive union coupling the column outlet with fused silica PicoTip emitter, ID 10 μm (New Objective, Inc.) and the ion transfer capillary temperature was set to 275°C. The mass spectrometer was operated in data-dependent mode using a full scan, m/z range 300–1,800, nominal resolution of 120,000, with AGC target set to standard with a maximum injection time of 50 ms and advanced peak detection enabled. Precursors were isolated in a range between 1 e4 and 1 e20 and MS/MS scans of the 40 most abundant ions were acquired in the Ion Trap in Turbo scan rate. MS/MS spectra were acquired by HCD fragmentation using normalized collision energy of 30%, isolation width of 1.6 m/z, resolution of 120,000, and a AGC target value set to standard and maximum injection time set to auto. Precursor ions with charge states 2-5 were selected for fragmentation and put on a dynamic exclusion list for 15 seconds. The peptide match feature was set to the preferred mode and the feature to exclude isotopes was enabled.

#### Phosphoproteomic data Processing and Peptide quantification

Raw data files were processed using Proteome discoverer 2.5 (Thermo Fisher Scientific) and searched against an in-house constructs and contaminants database and the Araport11 protein database. The processing workflow was made up of the Sequest HT search engine, Percolator (for target/decoy selection) and IMP-ptmRS (to calculate modification site probabilities). Tryptic peptides with up to 2 possible mis-cleavage and charge states +2, +3 were allowed in the search. The following peptide modifications were included in the search: carbamidomethylated Cysteine (fixed), oxidized Methionine (variable) and phosphorylated Serine, Threonine and Tyrosine (variable). Data were searched with a monoisotopic precursor and fragment ion mass tolerance 10 ppm and 0.6 Da respectively. Peptides were quantified using the ‘basic modification analysis’ consensus workflow provided by Proteome Discoverer 2.5 and expressed as abundance ratios. Peptides in the Peptide groups tab in the results files were filtered for ‘phospho’ and reliable and detectable ‘quan’ values. Threshold for differential phosphopeptides was set at minimum 2-fold change in abundance ratio and an adjusted abundance ratio p-value of less than 0.05. Data for Peptide groups were exported to Excel and processed in R.

#### Gene Ontology (GO) enrichment analysis

Gene Ontology (GO) enrichment analysis was performed in two steps. We first used the “GPHyperGParams” function in the R package “GOstats” to define the test parameters (PMID: 17098774). Biological process (BP) ontology was tested and a *P* value < 0.01 was used as the cut-off of enrichment significance. The Conditional parameter was set to TRUE to consider the structure of the GO graph when estimating each BP term. We second used the “hyperGTest” function in the same package to perform the hypergeometric test (PMID: 17098774). Then we displayed the top 10 (*P* value <0.01, Ranked by OddsRatio, Count ≥ 5 for PSR2 and Count ≥ 3 for PITG_15142) significantly enriched GO terms in the plot.

#### Statistical analysis

Data are represented as the mean ± s.e.m. or as box-and-whisker plots in which the centre line indicates the median, the bounds of the box indicate the upper and lower quantiles, using R Studio (https://www.r-project.org/) or GraphPad Prism 8.0. Statistical analyses performed using GraphPad Prism 8.0. Relative PP2A activities were analysed using a two-tailed Student t-test. Data for testing the significant differences between Arabidopsis genotypes were performed using One-way ANOVA and post hoc Tukey. Data and statistical analysis for MS analyses are provided in Table S4 and S5. Data and statistical analysis for additional analyses are provided in Table S6.

## SUPPLEMENTAL INFORMATION TITLES AND LEGENDS

TableS1. Data collection and refinement statistics of PDF1(1-390)-PSR2(59-670) and apo PITG_15142, related to Figure 2 and Figure 5

Table S2. The RMSD, sequence identity, and conservation score of key residues by comparing all WY-LWY motifs with PSR2(170-310), related to Figure 4

Table S3. Pairwise comparison of PP2A-interacting modules and C terminal LWY units, related to Figure 4

Table S4. Data lists of quantitative phosphoproteomic analysis by MS, related to Figure 5

Table S5. Significant candidates identified by coIP-MS using PSR2 and PITG_15142 as bait, related to Figure S5

Table S6. Statistical summary, related to Figure 1-2, 4 and Figure S1, S4 and S5.

Document S1. Structure prediction by AlphaFold2 and WYLWY unit arrangement of all LWY effectors in *P. infestans* and *P. sojae*, related to Figure 4.

