## Supplementary material for "Pathogen protein modularity enables elaborate mimicry of a host phosphatase": Table S1

**Table S1.** **Data collection and refinement statistics of PDF1(1-390)-PSR2(59-670) and apo PITG_15142, related to Figure 2 and 5**

|  | PDF1(1-390)-PSR2(59-670) | PITG_15142 |
| --- | --- | --- |
| **Data collection** |  |  |
| Space group | *P*2_1_2_1_2_1_ | *P*22_1_2_1_ |
| Cell dimensions |  |  |
| *a*, *b*, *c* (Å) | 76.21, 111.55, 161.76 | 47.85, 89.39, 129.37 |
| α, β, γ (°) | 90.00, 90.00, 90.00 | 90.00, 90.00, 90.00 |
| Resolution (Å) | 50.00-2.30 (2.38-2.30) * | 50.00-3.15 (3.26-3.15) * |
| *R*_sym_ or *R*_merge_ | 0.024 (0.64) | 0.065 (0.418) |
| *I* / σ*I* | 57.6 (3) | 20.5 (2.2) |
| Completeness (%) | 99.9 (99.6) | 99.1 (97.3) |
| Redundancy | 6.4 (6.5) | 4 (3.3) |
| **Refinement** |  |  |
| Resolution (Å) | 2.3 | 3.1 |
| No. reflections | 61,929 | 9,493 |
| *R*_work_ / *R*_free_ | 0.209/0.227 | 0.240/0.276 |
| No. atoms | 8,717 | 3,240 |
| Protein | 7,878 | 3,207 |
| Ligand/ion | 8 | 9 |
| Water | 831 | 24 |
| *B*-factors | 39.52 | 47.51 |
| Protein | 39.28 | 47.69 |
| Ligand/ion | N/A | 40.90 |
| Water | 41.49 | 25.40 |
| R.m.s. deviations |  |  |
| Bond lengths (Å)  Bond angles (°) | 0.005  0.799 | 0.004  0.912 |
| Ramachandran plot |  |  |
| Favored (%) | 98.28 | 90.80 |
| Allowed (%) | 1.72 | 8.71 |
| Outlier (%) | 0.10 | 0.5 |

*Values in parentheses are for highest-resolution shell.
