## Supplementary figures and images for "Pathogen protein modularity enables elaborate mimicry of a host phosphatase"

### Document S1

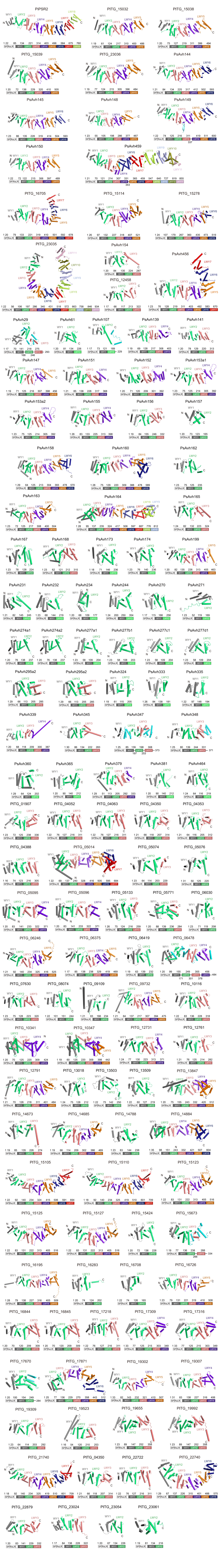
